# Multiomic profiling of a unique in-transit melanoma cohort identifies melanoma differentiation as predictor of tumor progression and therapy response

**DOI:** 10.1101/2025.10.01.679282

**Authors:** Giuseppe Tarantino, Anne Zaremba, Tuulia Vallius, Mark Woodnorth, Yichao Hua, Roxanne Pelletier, Mariana Lopez Leon, Yingxiao Shi, Zoltan Maliga, Samira Makhzami, Tyler J. Aprati, Bojan Karlaš, Valerie Glutsch, Bastian Schilling, Jessica Hassel, Carola Berking, Jochen Utikal, Friedegund Meier, Frank Meiss, Lucie Heinzerling, Katharina Kähler, Jiajia Chen, Lisa Zimmer, Antje Sucker, Elisabeth Livingstone, Eva Hadaschik, Christine Lian, George Murphy, Yevgeniy R Semenov, Genevieve M. Boland, Peter K. Sorger, Florian Rambow, David Liu, Dirk Schadendorf

## Abstract

Melanoma patients with in-transit metastasis (ITM), a stage of disease where melanoma has metastasized to sites in between the primary lesion and draining lymph node, vary significantly in their clinical outcomes, but the biology driving differential outcomes in ITM is poorly understood. To elucidate the mechanisms of differential outcomes, we utilized multimodal molecular profiling (WES, RNA-seq, highly multiplexed immunofluorescence, spatial transcriptomics) in 1) evolutionary analysis of longitudinal tumor samples and 2) identifying prognostic tumor intrinsic and microenvironmental features in a unique cohort of patients with unresectable ITM. Among other findings, we observed a persistent dedifferentiated AXL⁺/NGFR⁺ clonal lineage pre-existing and following immune checkpoint blockade in in-transit and distant metastases. Concordantly, we found that low pigmentation and high T cell exhaustion signatures were independently associated with distant progression. Our findings highlight tumor cell state and immune dysfunction as key predictors and potential biomarkers of metastatic risk in ITM.

**STATEMENT OF SIGNIFICANCE:** What drives distant progression in melanoma is unclear. Analyzing tumor and immune features in a rare in-transit melanoma patient cohort, we identify biological signals highlighting how immune and tumor states observable in pre-distant metastasis melanomas shape long-term outcomes, and nominate potential prognostic biomarkers.

## INTRODUCTION

Melanoma represents 5% of all skin cancers but is responsible for 95% of skin cancer-related deaths. Melanoma cells can spread to cause advanced metastatic disease in two ways: hematogenously and via the lymphatic system. Skin metastases near the primary tumor are considered an intralymphatic manifestation of melanoma^1,2^. While satellite metastases are defined as occurring within a 2 cm radius of the primary tumor, melanoma in-transit metastases (ITM) refer to lesions found between the primary melanoma and the nearest regional lymph node basin; ITM can develop in up to 20% of melanoma patients^3^. The presence of ITM has been linked to worse clinical outcomes compared to patients without ITM or satellite metastases^4^. In a significant subset of patients, locoregional recurrences occur until the disease becomes unresectable. However, clinical data also show a remarkably high rate of long-term survival without disease progression to distant sites in a subset of ITM patients^5^. The underlying mechanisms driving these distinct clinical outcomes remain unclear. Previous studies have shown that patients with ITM tend to be older than the average melanoma patient (>50 years), are more frequently female, and often have a primary melanoma located on the lower extremities, and involve both ulceration and greater Breslow thickness^2^ suggesting that ITM might have a specific biology^5^.

When ITM is confined to a regional area, resection can, in some cases, be a curative. However, if ITM becomes unresectable, both local and systemic treatment options must be explored. Isolated limb perfusion (ILP), electrochemotherapy, and intralesional therapy with agents including oncolytic viruses (e.g. TVEC), have shown varying degrees of response and are used either alone or in combination with immune checkpoint inhibitors (ICI)^6,7^. The literature is discordant with respect to the responsiveness of ITM to systemic therapy with ICI^8,9^. A Scandinavian group reported a favorable 56% response rate to anti-PD1 therapy alone in patients with in-transit metastases, with or without nodal involvement^10^. In contrast, other studies observed a lower overall response rate (ORR) of 31% to anti-PD1 in patients with unresectable ITMs^5^. One possible explanation for these differences in cohort composition is the variation in treatment approaches, specifically whether therapy was administered for unresectable ITM or unresectable nodal disease. Moreover, both studies were retrospective, and prospective data on treatment responses in ITM is still lacking. The lack of data is partly because patients with ITM were either excluded from or not clearly documented in pivotal clinical trials, often due to challenges in assessing response: RECIST measurements are difficult to apply to the small ITM lesions^10^.

The tumor and microenvironment of ITM have not been fully explored, and the role of ITM in melanoma progression, therapy response, and local treatment remains poorly understood. Preliminary findings from Bai et al. indicate that baseline samples from patients with ITM who had progressed after anti-PD1-based immunotherapy exhibited tumor heterogeneity and immune-exclusion characteristics^11,12^.

In this paper we characterize melanoma patients with unresectable stage III ITM and satellite metastasis for tumor characteristics, survival and therapy response. To our knowledge, this is the first cohort exclusively comprising melanoma patients initially characterized by the presence of unresectable ITM with multi-omics analysis. In addition, we analyzed multiple specimens from two such patients in detail using comprehensive multi-omic approaches to examine the dynamics and evolution of longitudinal tumor samples, aiming to identify the mechanisms driving these heterogeneous phenotypes. We integrated bulk sequencing (comprising both whole exome [WES] and RNA sequencing) and single-cell sequencing (at both RNA and protein levels) with spatial datasets to investigate heterogeneity and cell-type-specific progression pathways in melanoma patients with ITM undergoing systemic therapy. Matched tumor dissociates and formalin-fixed paraffin-embedded (FFPE) tissues as well as cryo-conserved melanoma lesions were processed for parallel sequencing and high-plex spatial profiling using cyclic immunofluorescence imaging (CyCIF)^13^. This made it possible to identify transcriptomic features associated with melanoma progression and spread to distant sites as compared to localized ITM disease. We also found patient-specific morphologies and mechanisms of resistance and aggressiveness in our longitudinal dataset characterizing melanoma ITM evolution on multiple layers.

## RESULTS

### Longitudinal Profiling of Two ITM Patient Trajectories Identifies Histopathological, Immune, and Transcriptomic Features Behind Diverging Outcomes

To investigate the molecular and morphological factors driving divergent outcomes in ITM patients, we utilized two complementary approaches: 1) longitudinal molecular profiling and evolutionary analysis based on whole exome sequencing (WES), bulk RNA-sequencing, single-nucleus RNA-sequencing, CyCIF imaging and spatial transcriptomics, of two patients with contrasting outcomes (i.e. treatment resistance and progressive disease vs. spontaneous remission) (**Figure 1A**); 2) analysis of pre-treatment tumor molecular features in a unique cohort of 41 ITM patients to evaluate tumor-intrinsic and microenvironmental features predicting subsequent distant progression (**Figure 1B**, **Table 1**).

**Figure 1.**
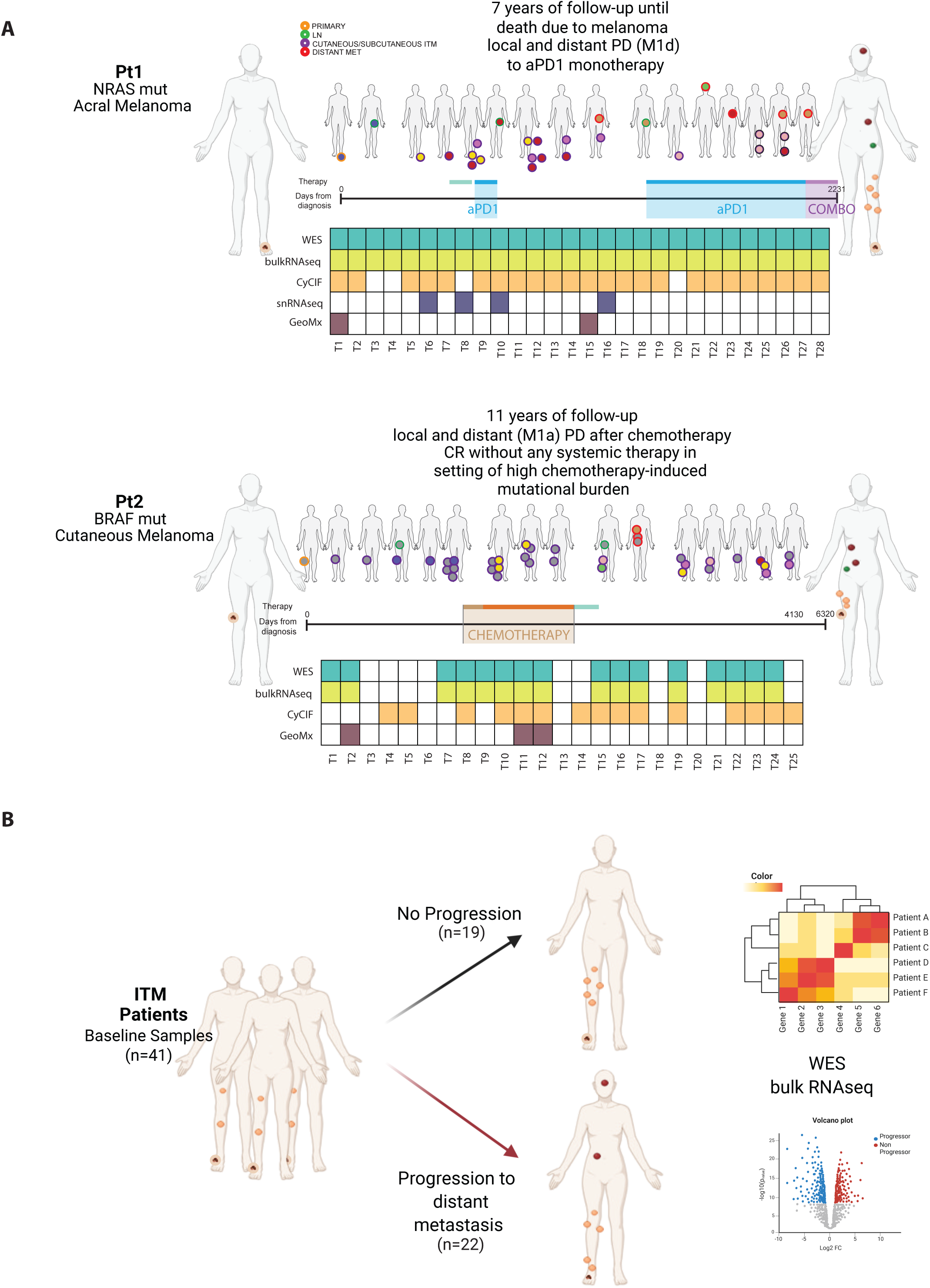
Longitudinal Multi-Modal Characterization and Baseline Analysis in Unresectable In-Transit Metastases (ITM) Melanoma Patients for Progression Assessment. **A.** Longitudinal multi-modal characterization of two melanoma patients. Patient 1 (NRAS-mutant acral melanoma) received single-agent anti-PD1 treatment with 7 years of follow-up and 28 biopsies collected. Whole exome sequencing (WES) and bulk RNA sequencing (RNAseq) were performed on all 28 biopsies, with additional characterization using cyclic immunofluorescence (CyCIF) on 24 samples, single-nucleus RNA sequencing (snRNAseq) on 4 samples, and GeoMX digital spatial profiling on 2 samples. The biopsy sites are depicted in the upper schematic representation. Patient 2 (BRAF-mutant cutaneous melanoma) underwent chemotherapy with 11 years of follow-up and 25 biopsies collected. WES and bulk RNAseq were performed on 16 samples, CyCIF on 15 samples, and GeoMX on 3 samples. Biopsy locations are shown in the lower schematic representation. **B.** Baseline analysis of a cohort of 41 patients with unresectable in-transit metastases (ITM). Baseline ITM biopsies were evaluated for whole exome sequencing (WES) and bulk RNA sequencing (RNAseq) data. Patients were stratified into those who progressed to distant metastasis and those who did not.

**Table 1.** Clinical Characteristics and sequenced samples of the unresectable ITM patients evaluated.

| Characteristic | N | Overall, N = 41<sup>†</sup> | Distant Progressors, N = 22<sup>†</sup> | Distant non-progressors, N = 19<sup>†</sup> |
| --- | --- | --- | --- | --- |
| **Sex** | 41 |  |  |  |
| female |  | 17 (41.5%) | 10 (45.5%) | 7 (36.8%) |
| male |  | 24 (58.5%) | 12 (54.5%) | 12 (63.2%) |
| **Age at diagnosis** | 41 | 67 (12) | 63 (13) | 72 (9) |
| **Localisation primary melanoma** | 41 |  |  |  |
| acral |  | 6 (14.6%) | 4 (18.2%) | 2 (10.5%) |
| head_neck |  | 2 (4.9%) | 0 (0.0%) | 2 (10.5%) |
| lower_extremity |  | 22 (53.7%) | 13 (59.1%) | 9 (47.4%) |
| other |  | 1 (2.4%) | 1 (4.5%) | 0 (0.0%) |
| trunk |  | 6 (14.6%) | 2 (9.1%) | 4 (21.1%) |
| upper_extremity |  | 4 (9.8%) | 2 (9.1%) | 2 (10.5%) |
| **Histological subtype** | 41 |  |  |  |
| ALM |  | 8 (19.5%) | 5 (22.7%) | 3 (15.8%) |
| amelanotic |  | 2 (4.9%) | 2 (9.1%) | 0 (0.0%) |
| LMM |  | 1 (2.4%) | 0 (0.0%) | 1 (5.3%) |
| NMM |  | 18 (43.9%) | 8 (36.4%) | 10 (52.6%) |
| other |  | 1 (2.4%) | 1 (4.5%) | 0 (0.0%) |
| SSM |  | 4 (9.8%) | 2 (9.1%) | 2 (10.5%) |
| unknown |  | 7 (17.1%) | 4 (18.2%) | 3 (15.8%) |
| **Breslow thickness** | 41 |  |  |  |
| T1 |  | 2 (4.9%) | 0 (0.0%) | 2 (10.5%) |
| T2 |  | 4 (9.8%) | 3 (13.6%) | 1 (5.3%) |
| T3 |  | 17 (41.5%) | 9 (40.9%) | 8 (42.1%) |
| T4 |  | 17 (41.5%) | 9 (40.9%) | 8 (42.1%) |
| unknown |  | 1 (2.4%) | 1 (4.5%) | 0 (0.0%) |
| **Ulceration** | 41 |  |  |  |
| no |  | 13 (31.7%) | 9 (40.9%) | 4 (21.1%) |
| unknown |  | 7 (17.1%) | 1 (4.5%) | 6 (31.6%) |
| yes |  | 21 (51.2%) | 12 (54.5%) | 9 (47.4%) |
| **Systemic_therapy** | 41 | 30 (73.2%) | 14 (63.6%) | 16 (84.2%) |
| **Firstline PD-1 yes/no** | 41 | 16 (39.0%) | 5 (22.7%) | 11 (57.9%) |
| **BOR to antiPD-1** | 16 |  |  |  |
| PD |  | 6 (37.5%) | 2 (40.0%) | 4 (36.4%) |
| PR |  | 4 (25.0%) | 1 (20.0%) | 3 (27.3%) |
| SD |  | 5 (31.3%) | 2 (40.0%) | 3 (27.3%) |
| UNK |  | 1 (6.3%) | 0 (0.0%) | 1 (9.1%) |
| **WES** | 41 |  |  |  |
| N |  | 4 (9.8%) | 2 (9.1%) | 2 (10.5%) |
| Y |  | 37 (90.2%) | 20 (90.9%) | 17 (89.5%) |
| **RNAseq** | 41 |  |  |  |
| N |  | 5 (12.2%) | 0 (0.0%) | 5 (26.3%) |
| Y |  | 36 (87.8%) | 22 (100.0%) | 14 (73.7%) |
| <sup>†</sup> n (%); Mean (SD) |  |  |  |  |

The first patient (Pt1) a 59-year-old female, was diagnosed in 2013 with an ulcerated acral lentiginous melanoma, Breslow thickness 4.6 mm, located on the left sole with a NRAS Q61H mutation. Initial treatment included wide local excision and sentinel lymph node biopsy, which revealed metastasis in a left inguinal lymph node (primary tumor stage IIIB pT4b pN1a M0). One year later, the patient developed a recurrence with a left inguinal lymph node metastasis. Adjuvant interferon-gamma therapy was declined. Imaging showed no further metastases, and a lymphadenectomy was performed, revealing a 2.2 cm lymph node metastasis without evidence of extracapsular extension. Four months post-lymphadenectomy, the patient developed the first ITM in the left plantar region, which was surgically excised. Over the subsequent six months, multiple additional ITMs emerged on the left foot, lower leg, and thigh, rendering the disease unresectable. PET-CT imaging revealed bilateral pulmonary nodules and confirmed the presence of multiple in-transit and satellite metastases on the plantar foot and left leg.

Pt1 was enrolled in a phase Ib/II clinical trial investigating LEE011 (CDK4/6 inhibitor) and MEK162 (MEK inhibitor) for patients with NRAS-mutant melanoma. Treatment was discontinued after four weeks due to patient preference in the context of grade 1 toxicity. Multiple resections of ITMs were performed both prior to and following systemic therapy.

Subsequently, anti-PD-1 therapy with pembrolizumab was initiated but was paused after a single infusion due to the onset of autoimmune thyroiditis. Progressive locoregional disease necessitated repeated surgical interventions on the left foot and leg, including resection of a 7 cm left iliac lymph node metastasis, which showed no capsular involvement.

The patient opted against further systemic therapy and declined isolated limb perfusion and underwent multiple additional surgical resections instead. Twelve months later, progression in the left iliac/inguinal region required further surgical intervention, which resulted in an R1 resection margin. Anti-PD-1 therapy with pembrolizumab was resumed. Follow-up imaging after four treatment cycles demonstrated progression in both the lungs and left leg skin. Therapy escalation was advised but declined; pembrolizumab monotherapy was continued. Six months into treatment, imaging revealed cerebellar metastases (stage M1d) along with further systemic disease progression. Cerebral surgery was recommended for symptomatic brain metastases, and therapeutic intensification discussed. The patient chose to continue anti-PD-1 therapy beyond progression and declined brain radiotherapy. Additional surgical procedures were carried out to manage locoregional cutaneous progression on the left leg and lower abdomen.

Subsequent imaging revealed gastric wall thickening and diagnostic work-up confirmed the presence of a gastric melanoma metastasis, alongside newly developed ITMs on the left leg. Clinical trial enrollment was offered but declined by the patient and pembrolizumab therapy was continued beyond oligo-progression (bi-pulmonary, gastric, and distant cutaneous metastases). With further disease progression, therapy was escalated to combined ICI with ipilimumab and nivolumab. After one cycle, treatment was de-escalated to nivolumab monotherapy due to patient preference. Systemic therapy was ultimately discontinued, and the patient transitioned to best supportive care. The patient died seven years after the initial melanoma diagnosis due to progressive metastatic melanoma.

Over the period of 7 years, Pt1 developed several ITMs (n=19 available longitudinal tumor biopsies) and several distant metastases (n=5 available longitudinal tumor biopsies), including a brain (T22) and gastric (T27 and T28) metastases.

The second patient (Pt2), a 67-year-old female, had a BRAF V600E^mut^ primary cutaneous melanoma located on the right knee. Two years following initial surgical treatment, the patient developed a local recurrence, which was managed with a wide local excision. Six months later, multiple cutaneous satellite metastases appeared, and a biopsy of the left inguinal lymph nodes confirmed melanoma lymph node metastasis. The disease showed locoregional progression with further cutaneous metastases, necessitating multiple surgical excisions. Systemic therapy was initiated as part of a clinical trial with dacarbazine and PTNZK, followed by dacarbazine monotherapy. Despite systemic therapy, the patient continued to develop cutaneous and subcutaneous metastases on the right upper and lower leg, which were managed surgically. Due to ongoing progression, isolated limb perfusion (ILP) of the right leg was performed using tumor necrosis factor-alpha (TNF-α) and melphalan. This was followed by surgical excision of residual locoregional disease, including bilateral inguinal lymph node and subcutaneous metastases including M1a subcutaneous metastasis. Over the subsequent six years, the patient underwent multiple local excisions for recurrent in-transit metastases. The final in-transit metastasis was surgically removed 11 years after the initial melanoma diagnosis. Over the course of 11 years, Pt2 developed numerous ITMs, and distant cutaneous and subcutaneous metastases, yet remained in complete remission thereafter without any additional systemic therapy and is still alive 9 years later (06/2023).

Histopathological evaluation of H&E-stained sections identified distinct morphological features for Pt1 and Pt2 tumors. Hallmarks of aggressive growth (clefting, ulceration, and poorly differentiated morphology) were found in the primary tumor from Pt1 (T1, **Figure 2A**). In contrast, the first available tumor from Pt2 (ITM) exhibited morphological features resembling inflammatory regression; these included fibrosis and abundant immune infiltration following chemotherapy, suggestive of an effective anti-tumor immune response (**Figure 2B**). Immune composition, as assessed by CyCIF analysis, revealed profound differences in the composition of tumor microenvironment (TME) in Pt1 and Pt2. Pt1 exhibited a low level and heterogeneous immune infiltrate, with the immune component of the TME predominantly composed of CD163+ macrophages, which are often associated with immune suppression (**Figure 2C&E**). In contrast, Pt2 demonstrated a significantly higher immune infiltrate, including a high abundance of CD8+ T cells within the tumor and at the invasive boundary (**Figure 2D&F**). These CD8+ T cells expressed GZMB, implying cytotoxicity (**Supplementary Figure 1A&B**). CyCIF & Bulk RNAseq showed low expression of AXL and NGFR in pt2 samples (**Figure 2** & **Supplementary Figure 1D**). While there was significant intertumoral heterogeneity in signatures of melanocytic vs dedifferentiation in both patients in bulk RNAseq (**Supplementary Figure 1C&D**), tumors from Pt1 had higher proportions of tumor cells expressing NGFR or AXL, markers of neural crest and dedifferentiated tumor states (**Figure 2G&H**). Notably, across tumors, we found good correlation between immune infiltration and tumor state inferred from bulk RNAseq vs CyCIF (T16 and T25 from Pt1, **Supplemental Figure 2**).

**Figure 2.**
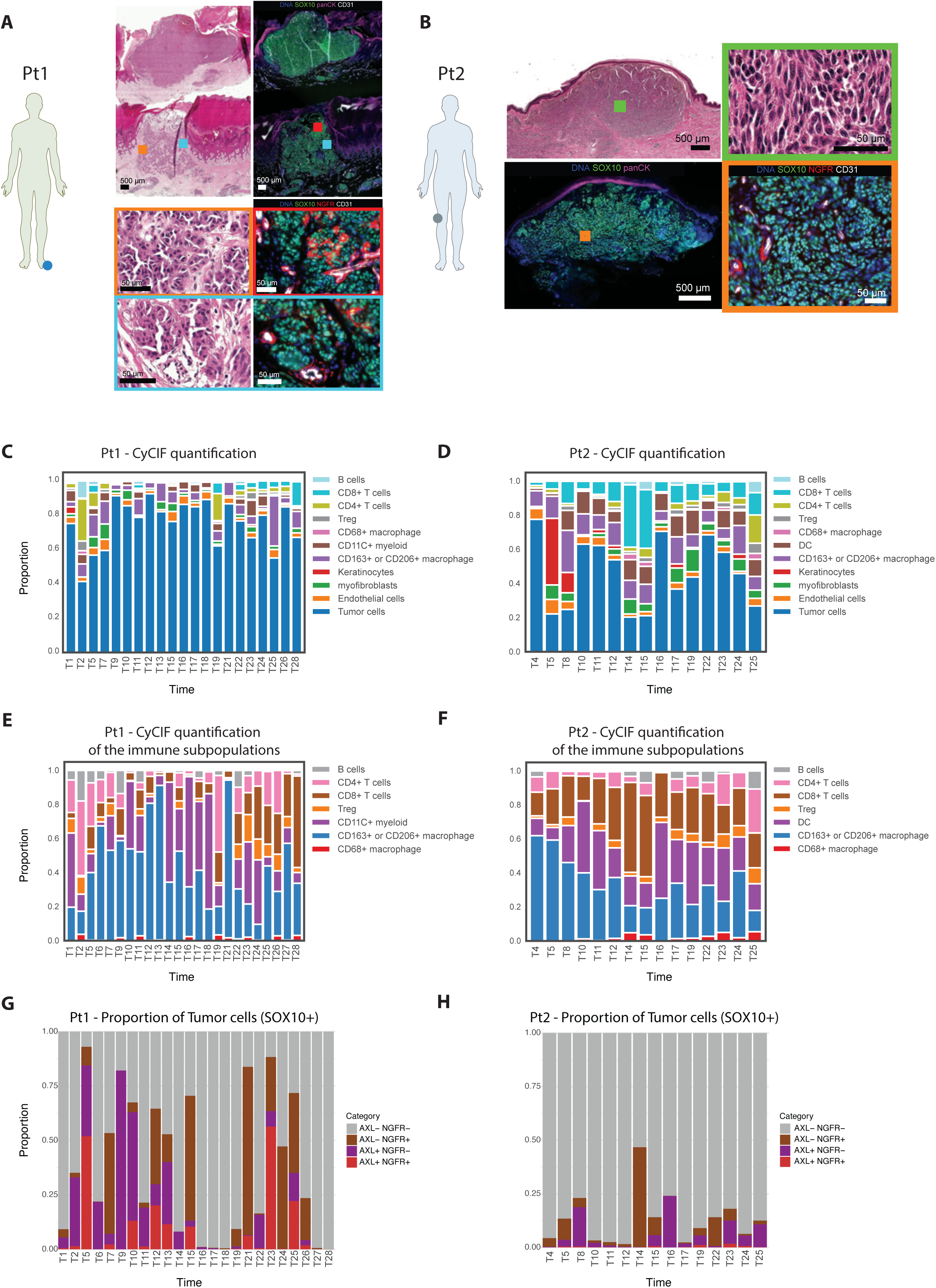
Histopathological Tumor and Immune Profiling of Primary, Unresectable In-Transit, and distant metastasis Melanoma Tumors in Pt1 and Pt2. **A.** Histopathological and CyCIF images of the primary tumor sample from Patient 1 (Pt1) displaying high histopathological heterogeneity and aggressive growth features such as clefting and ulceration. **B.** H&E and CyCIF images of the baseline in-transit metastasis (ITM) sample from Patient 2 (Pt2) showing lower histopathological heterogeneity and reduced clefting compared to Pt1. **C.** CyCIF profiling of Pt1 samples as stacked bar plot of the general cellular compartments. **D.** CyCIF profiling of Pt1 samples as stacked bar plot of the general cellular compartments. **E.** CyCIF Immune profiling showing the major immune subpopulations of Pt1. **F.** CyCIF Immune profiling showing the major immune subpopulations of Pt2. **G.** CyCIF quantification of SOX10^+^ cells in Pt1, showing in red the proportion of cells AXL^+^ and NGFR^+^, in purple the AXL^+^ NGFR^−^, in brown AXL^−^ NGFR^+^ and in gray AXL^−^NGFR^−^. **H.** CyCIF quantification of SOX10^+^ cells in Pt2, showing in red the proportion of cells AXL^+^ and NGFR^+^, in purple the AXL^+^ NGFR^−^, in brown AXL^−^ NGFR^+^ and in gray AXL^−^NGFR^−^.

### Resistance-Associated Mutations in Pt1 Lineages and the Persistence of a Single Lineage After ICI Treatment

Longitudinal whole-exome sequencing (WES) data was used for phylogenetic characterization of ITM tumor samples from Pt1 and Pt2. In common with other acral samples in our cohort, Pt1 tumors had a high genomic heterogeneity, as defined by the proportion of subclonal mutations divided by the total nonsynonymous mutations. Consistent with this, across all Pt1 tumors from this patient we observed 7 distinct tumor lineages, which existed prior to treatment and evolved over the course of the disease (**Figure 3A**). Pt1-L1 comprised the common ancestral and earliest detected clones (found in the primary tumor and a draining lymph node) from which remaining lineages were derived. Pt1-L3 and L4 arose from a shared ancestral clone, were detected early, and manifested only as in-transit lesions. Pt1-L2 arose from the same shared ancestral clone as Pt1-L3 and Pt1-L4, but split from the other lineages early (with acquisition of a clonal NF1 loss of function mutation) and presented 4 years later as a brain metastasis. This metastasis exhibited the highest tumor mutational burden (TMB) and degree of aneuploidy among all metastases, suggesting that genetic isolation and selective pressures within the brain microenvironment shaped its genomic profile. Pt1-L3 and -L4 only were detected early and disappeared shortly following treatment with anti-PD1 whereas “late-stage” lineages (Pt1-L5, -L6, and -L7) were detected years after aPD-1 therapy, included distant metastases, and displayed distinct molecular features. In Pt1-L7 these features included mutations in CUX1, FBXO11 and ARID2, reflecting a potential role in immune evasion or therapy resistance^14,15^. Pt1-L6 comprised most distant metastases and was characterized by high levels of aneuploidy (driven by the distant metastasis, **Supplementary Figure 3**), elevated TMB, and a likely pathogenic missense mutation in TET2. Pt1-L5 was the only lineage detected in both in-transit and distant metastases and was also the only lineage present before treatment and after treatment with anti-PD1. No specific resistance-associated mutations were identified in Pt1-L5, but the persistence of this lineage across disease stages highlights its metastatic potential.

**Figure 3.**
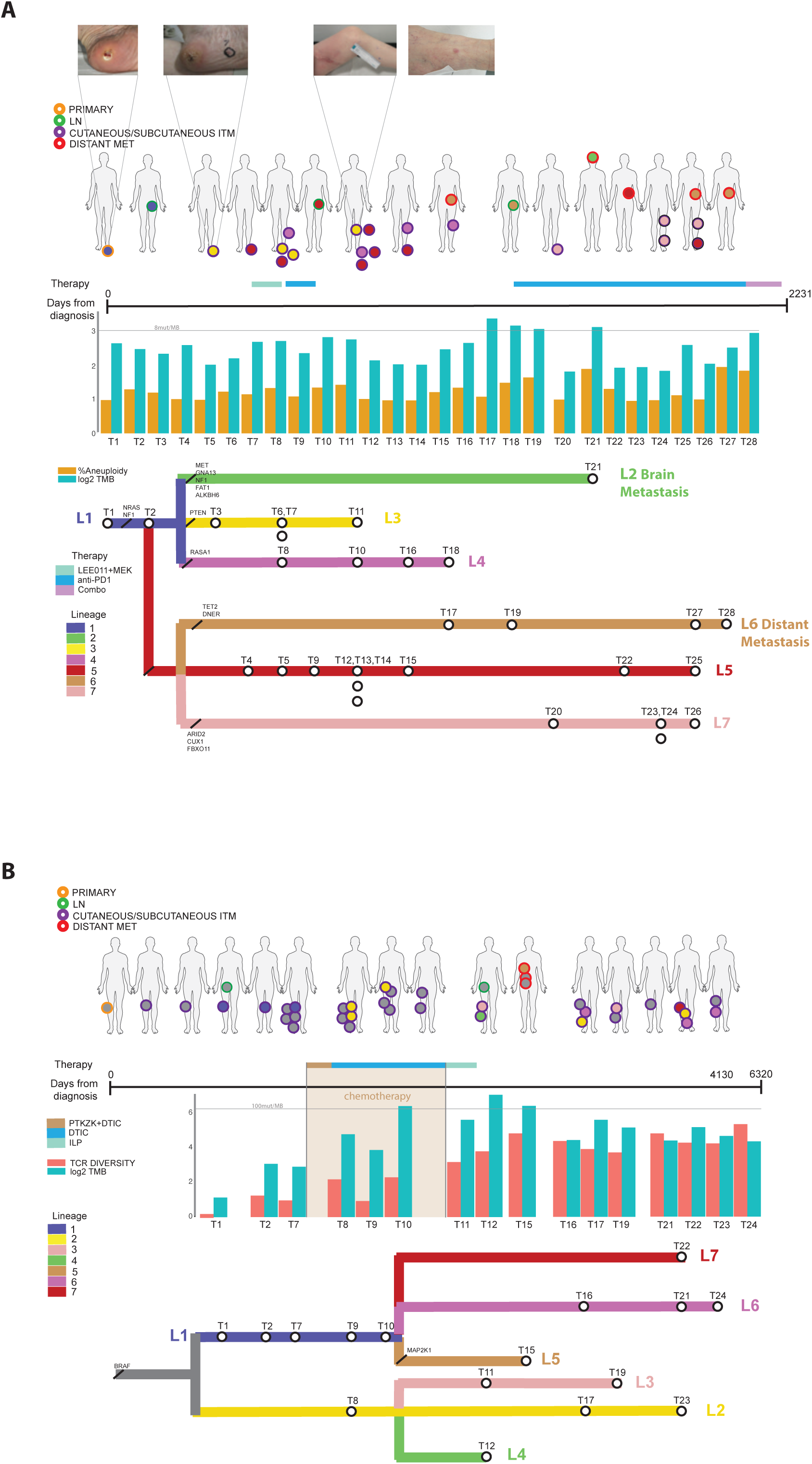
Evolution of Metastatic Lineages in Unresectable In-Transit Melanoma Patients: Clinical Progression and Treatment Context. **A.** Genomic lineage reconstruction in Patient 1 (Pt1). Biopsy sites are highlighted at the top of the plot, with treatment timeline shown above the x-axis (Days from diagnosis). The bar plot displays the percentage of aneuploidy (gold) and log2-transformed tumor mutational burden (TMB, cyan). Distinct metastatic lineages are represented in different colors, with associated mutations indicated in black. **B.** Genomic lineage reconstruction in Patient 2 (Pt2). Biopsy sites are highlighted at the top of the plot, with the treatment timeline represented above the x-axis (Days from diagnosis). The bar plot shows T-cell receptor (TCR) diversity estimated using TRUST4 (red) and log2-transformed TMB (cyan). Lineages are represented in different colors, with associated mutations indicated in black.

### Autonomous and effective host-immune response leading to spontaneous complete remission in a chemotherapy-induced high tumor mutational burden setting

Pt2 was diagnosed with cutaneous melanoma, characterized by a clonal BRAF V600E mutation, and developed distant stage M1a metastases as well as multiple ITMs that in aggregate represented seven co-evolving lineages (**Figure 3B**). Among these, only Pt2-L5 was associated with distant metastatic lesions and characterized by an acquired MAP2K1 mutation (in addition to the BRAF mutation in the common ancestor) suggesting its role in disease progression. The genomic evolution of tumors in Pt2 revealed a dynamic shift in mutational signatures over the course of the disease. The earliest three tumor samples (all ITMs) were enriched in a UV mutational signature, consistent with the cutaneous origin of the melanoma. However, following exposure to dacarbazine chemotherapy, subsequent tumors had dramatically higher TMB, which rose from 10 mutations/megabase (mut/MB) initially to >100 mut/MB following chemotherapy. Mutational signature analysis showed predominance of COSMIC mutational signature 11, which is associated with exposure to alkylating agents^16,17^ (**Supplementary Figure 4**). The increase in TMB was paralleled by an expansion in T cell receptor (TCR) diversity as estimated from bulk RNAseq, consistent with enhanced immunogenicity of the tumor arising from TMB-associated neoantigen generation^18^ (**Figure 3B**). CyCIF immunoprofiling revealed abundant Ki67+ and GZMB+ CD8+ tumor-infiltrating lymphocytes (TILs), suggestive of an active cytotoxic T cell response. Histopathological evaluation of tumor samples also demonstrated regression-like stromal changes, further supporting the hypothesis of an effective tumor-immune response was present (**Supplementary Figure 5**). Together, we postulate that the presence of high chemotherapy-induced TMB, increased TCR diversity, and presence of infiltrating CD8+ GZMB+ T cells contributed to a strong anti-tumor immune response that ultimately led to a disease remission and no further requirement for systemic therapy even in the presence of remaining unresectable ITMs on the trunk.

### Pigmentation Signature as a Predictor of Non-Progression to Distant Metastasis in ITM Patients

Building on the insights of Pt1 and Pt2, where divergent outcomes were influenced by genomic heterogeneity, tumor mutational burden, and immune responses, we profiled a cohort of single specimens from 41 ITM patients to identify features associated with progression or non-progression to distant metastasis. This cohort, consisting of ITM samples that had developed prior to the exposure of any systemic melanoma treatment (treatment-naïve) included 19 non-progressing patients and 22 patients who developed distant melanoma metastases (**Figure 1B**; **Table 1**), allowing for a comparative analysis of molecular and transcriptomic profiles. As a first step we compared the somatic mutations in the two groups of patients, and found no evidence for enrichment of any single event (after multiple hypothesis correction; **Supplementary Figure 6**). However, non-progressors were enriched for signatures associated with melanocytic tumor cell states (**Figure 4A&B**). In particular, the pigmentation signature defined by Rambow et al. (2018) emerged as the most significant feature distinguishing non-progressors (M0) from progressors (M1 disease). This signature reflects a differentiated tumor state in which pigmentation-related genes are upregulated, yet tumor cells are still proliferative (Ki67^+^,PCNA^+^). In our cohort this signature was associated with improved distant metastasis-free survival (DMFS; **Figure 4C**). To further investigate transcriptional tumor states in ITM, we evaluated a collection of signatures from Hu et al. 2024^19^ and identified additional signatures that stratify progressors and non-progressors. The top two signatures were identified by Rambow et al 2018 and Belote et al. 2021^20,21^. Interestingly, there was no overlap in genes between the Rambow pigmentation signature and the Belote Cmel signature (**Supplementary Figure 7A**); however, Gene Set Enrichment Analysis (GSEA) showed that the upstream regulators of both signatures are the same, involving MITF as a key transcriptional driver. Among the top differentially expressed genes, CAV1, a gene previously associated with decreased metastatic potential, was enriched in non-progressor samples^22^. Acral ITM patients were similar to Pt1 in exhibiting the highest enrichment of differentiated signatures and the highest genomic heterogeneity (**Supplementary Figure 7B**). This finding parallels the high genomic heterogeneity observed in Pt1 and suggests that the interplay between differentiation trajectory and genomic instability may contribute to the unique metastatic patterns seen in acral melanoma. To ensure that the observed associations were not confounded by microenvironmental signals, we also performed bulk RNA-seq deconvolution to isolate the tumor cell component and re-evaluated the enrichment of key signatures. Notably, the pigmentation signature remained significantly associated with non-progression (p = 0.014) even when assessed solely within the tumor compartment, whereas the Belote Cmel signature demonstrated a trend (p = 0.053) (**Supplementary Figure 7C**).

**Figure 4.**
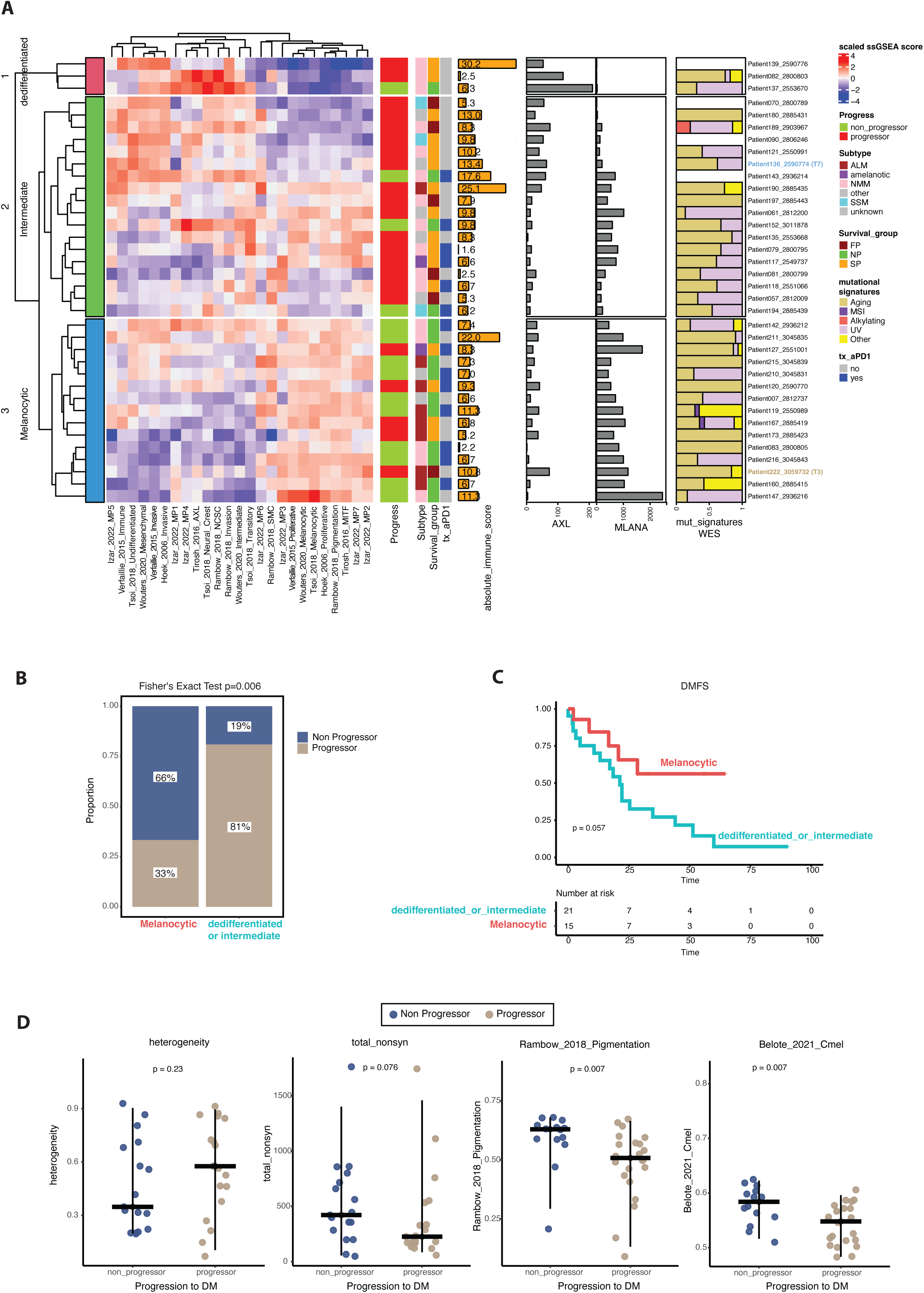
Melanoma Differentiation Status Predicts Metastatic Progression. **A.** Heatmap displaying the clustering of patients based on bulk RNA sequencing data. Patients who progressed to distant metastasis are marked in red. Clinical features (Subtype, Survival Group, Treatment) and bulk RNA sequencing features (Absolute Immune Score, AXL expression, MLANA expression) alongside whole exome sequencing (WES) features (mutational signatures) are shown on the right. **B.** Stacked bar plot showing the proportion of progressors and non-progressors stratified by melanocytic and dedifferentiated/intermediate groups. **C.** Distant metastasis-free survival (DMFS) comparing the melanocytic group versus the dedifferentiated/intermediate group. **D.** Genomic features (tumor heterogeneity and tumor mutational burden, TMB) and bulk RNA sequencing signatures (Rambow pigmentation signature and Belote cmel signature) comparing progressors and non-progressors. Statistical comparisons and p-values are indicated.

### Enrichment of T Cell Exhaustion Signatures in Progressors and Their Independent Predictive Value from the Pigmentation Signature

To investigate immune-related mechanisms associated with disease progression, we examined immune signatures from Mao et al.^23^ that were derived from a meta-analysis of gene expression signatures from immunotherapy-treated melanoma patients. These signatures are of particular interest due to their availability as targeted assays, making them directly applicable in clinical practice. Among these, the Exhausted_CD8_Nanostring signature was significantly enriched in patients with progressive disease, suggesting a potential role for T cell exhaustion in metastatic progression and treatment response. Based on this finding, we expanded our analysis to include additional exhaustion-related signatures, including those from Tirosh et al., derived from scRNASeq of metastatic melanomas^24^. Notably, all of the exhaustion signatures we evaluated were enriched in progressor samples (**Figure 5A**), supporting the hypothesis that immune dysfunction contributes to disease progression. To refine this observation, we assessed the top differentially expressed genes from these exhaustion signatures and identified EOMES, a well-established marker of T cell exhaustion^25^, as the gene with the highest fold-change between progressors and non-progressors (**Figure 5B**). Among all signatures evaluated *Tirosh CD4 T cells exhaustion* demonstrated the strongest stratification between progressors and non-progressors.

**Figure 5.**
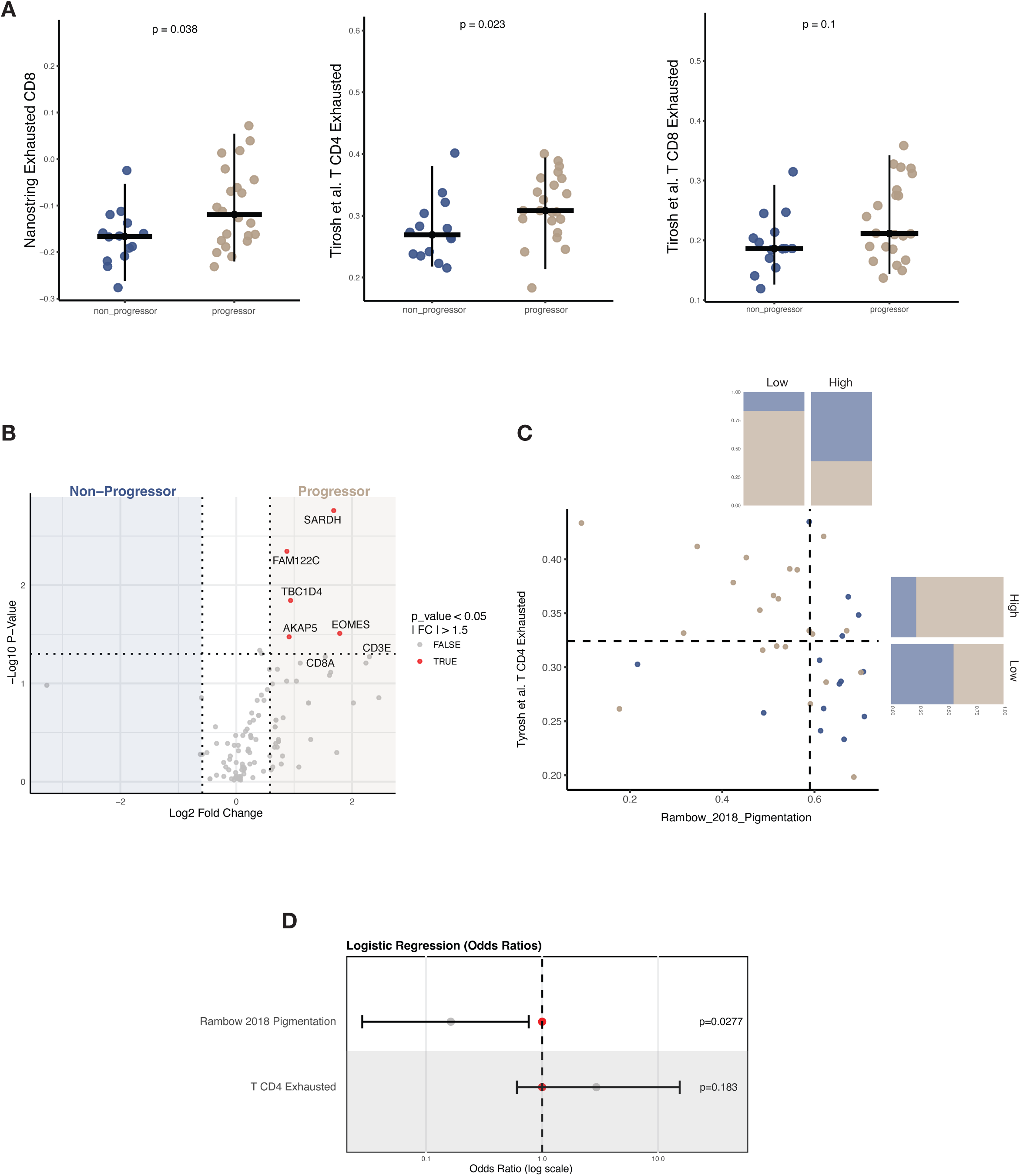
T cells exhaustions markers association with distant metastasis. **A.** Box plots comparing exhaustion signature scores between non-progressors and progressors for three immune exhaustion signatures: *NanoString Exhausted CD8* (left), *Tirosh et al. T CD4 Exhausted* (middle), and *Tirosh et al. T CD8 Exhausted* (right). The *NanoString Exhausted CD8* and *Tirosh T CD4 Exhausted* signatures were significantly enriched in progressors (*p* = 0.038 and *p* = 0.023, respectively), while *Tirosh T CD8 Exhausted* showed a non-significant trend (*p* = 0.1). **B.** Volcano plot displaying the top differentially expressed genes from exhaustion-related signatures between non-progressors and progressors. The exhaustion marker *EOMES* exhibited the highest fold-change in progressors (highlighted in red), along with other genes associated with T cell dysfunction. **C.** Scatter plot showing the relationship between the *T CD4 Exhausted* signature and the *Rambow 2018 Pigmentation* signature. Patients were stratified using a median split for each signature, revealing four distinct groups. Bar plots indicate the proportion of patients in each category, highlighting an inverse relationship between pigmentation and exhaustion. **D.** Logistic regression model demonstrating that the *Rambow 2018 Pigmentation* signature and the *T CD4 Exhausted* signature are independent predictors of disease progression. The pigmentation signature shows a protective effect with a significant odds ratio (*p* = 0.0277), while the exhaustion signature is associated with a higher risk of progression but does not reach statistical significance (*p* = 0.183).

To better understand the relationship between tumor differentiation and immune exhaustion, we examined the Tirosh TCD4 exhausted signature in conjunction with the pigmentation signature, splitting the cohort into 4 subgroups based on high and low levels of these two signatures split at the median (**Figure 5C**). Multivariable logistic regression revealed that the exhaustion and pigmentation signatures independently predicted disease progression. The pigmentation signature was associated with a protective effect, with a statistically significant and favorable odds ratio, while the T CD4 exhausted signature emerged as a strong risk factor with a high odds ratio (**Figure 5D**).

### Association of aPD1 treatment with decreased distant progression

Treatment with anti-PD1 was significantly associated with non-progression to distant metastasis (Fisher Exact Test p =0.029, OR=0.22; **Supplementary Figure 8A**). Strikingly, this was discordant with local response to aPD1, where response was similar between progressors and non-progressors (total n=16 pts treated with aPD1; local responses n= 4 (PR), n=5 (SD), n=6 (PD), n=1 (unknown)), **Supplementary Figure 8B**). This suggests that despite a lack of difference in local aPD1 response rates, there may be a systemic effect of aPD1 treatment with decreased distant progression. Indeed, we saw that 3/6 patients who had locally progressive disease as best response to aPD1 did not develop distant metastasis (**Supplementary Figure 8B**).

Importantly, when the pigmentation signature was incorporated into a multivariate model that also included anti-PD1 treatment status, pigmentation remained a significant and independent predictor of disease progression (**Supplementary Figure 9**). This suggests that tumor differentiation status, as captured by the pigmentation signature, plays a critical role in shaping metastatic potential, irrespective of exposure to immune checkpoint blockade.

### Role of dedifferentiated, AXLhigh/NGFRhigh Cells in Driving Distant Metastasis

To further investigate drivers of distant metastasis, we examined longitudinal samples from Pt1 and Pt2, focusing on the interplay between pigmentation-signature activities and the presence of AXL+ and NGFR+ cells (**Figure 6**). In Pt1, strikingly, Pt1-L5 had the lowest pigmentation-signature scores among all the lineages, and was also the only lineage detected across ITMs, distant metastases, and before and after treatment, suggesting a link between a dedifferentiated state that aligns with aggressive behavior (**Figure 6A**). Using CyCIF, we identified protein markers that correlated significantly with an RNA pigmentation signature; these comprise cells that were classified as staining positive for MART1, Ki67, PCNA, and, to a lesser extent, PMEL (**Supplementary Figure 9**). This suggests that the pigmentation signature captures a proliferative and differentiated melanoma cell state, the low enrichment of this signature in Pt1-L5 reinforce its role as a key factor in disease stratification. CyCIF analysis further demonstrated that L5 had the highest proportion of AXL+ and NGFR+ cells among all lineages, emphasizing the role of neural crest lineages in driving metastatic progression (**Figure 6C**). In Pt2, the only distant metastasis that was present, also displayed the lowest pigmentation signature score among all samples, reinforcing the association between low pigmentation scores, dedifferentiation, and metastatic potential (**Figure 6B**).

**Figure 6.**
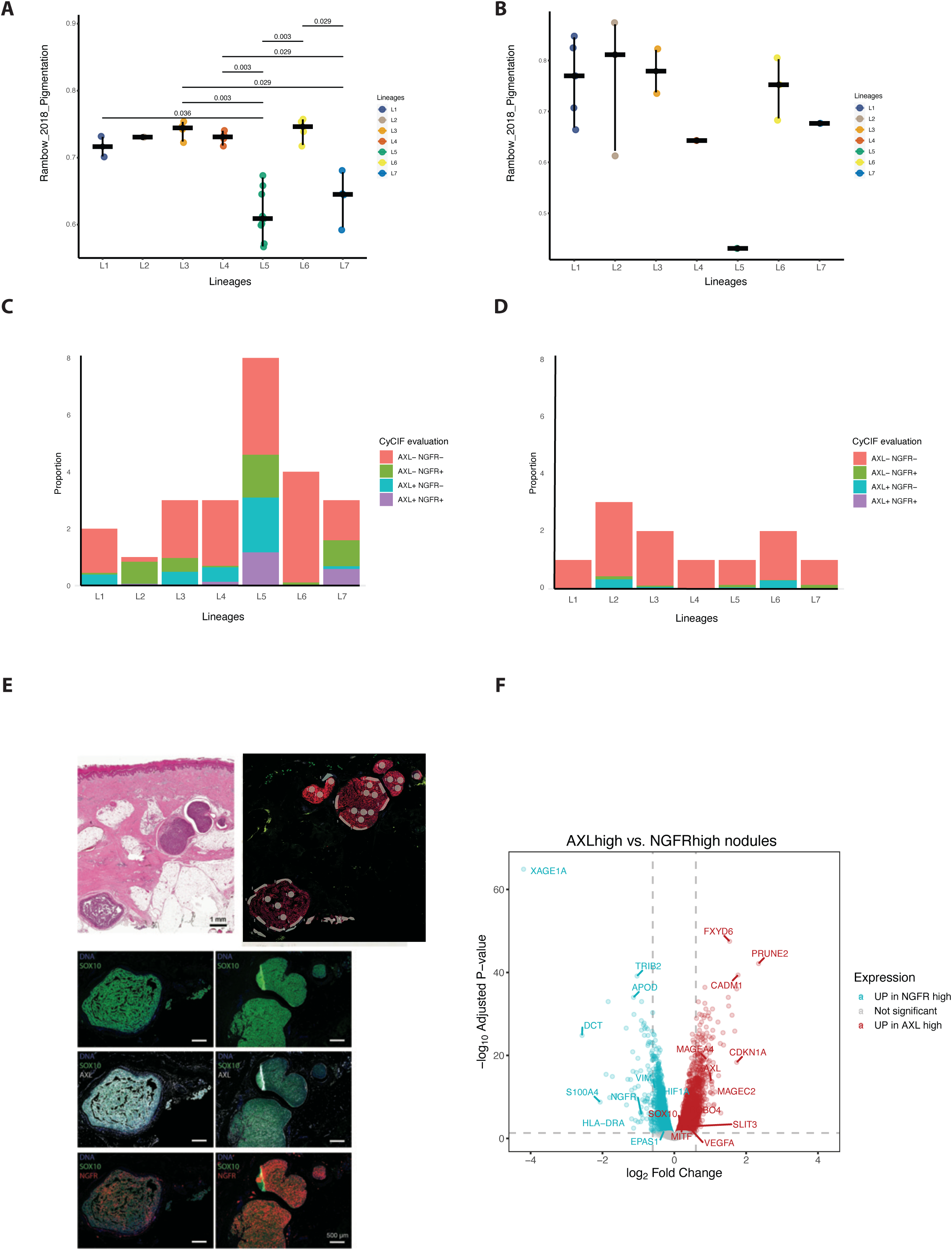
Pigmentation Signature, AXL, and NGFR Expression in Pt1 and Pt2: Lineage-Specific Insights into Metastasis. **A.** Box plot showing the pigmentation signature values estimated from bulk RNA sequencing across different genomic lineages in Patient 1 (Pt1). **B.** Box plot showing the pigmentation signature values estimated from bulk RNA sequencing across different genomic lineages in Patient 2 (Pt2). **C.** Evaluation of NGFR and AXL positive cells using CyCIF across the different lineages of Pt1. **D.** Evaluation of NGFR and AXL positive cells using CyCIF across the different lineages of Pt2. **E.** Histological and multiplex imaging of sample T15 from Pt1, showing regions of interest (ROIs) characterized with GeoMX spatial profiling. **F.** Volcano plot comparing nodules with high NGFR expression (cyan) and high AXL expression (red), highlighting differentially expressed genes between these two groups.

To delve deeper into the phenotypic heterogeneity of L5 in Pt1, we analyzed a highly heterogeneous in-transit metastasis sample (T15), consisting of three separate micronodules, using GeoMx digital spatial profiling (**Figure 6E**). CyCIF revealed that T15 contained two distinct nodules, one characterized by high AXL expression and the other by high NGFR expression. GeoMx data allowed us to select multiple regions of interest (ROIs) from each nodule for differential expression analysis. The differential expression analysis identified distinct melanoma antigens uniquely expressed in each nodule. In the NGFR-high nodule, XAGE1A, DCT, and S100A4 were significantly overexpressed, while the AXL-high nodule showed overexpression of MAGEA4 and MAGEC2 (**Figure 6F**). These findings underscore the phenotypic and functional diversity within Pt1-L5 and suggest that specific subpopulations may contribute uniquely to its metastatic behavior.

### Association of Pigmentation Signature Decrease with ICB Treatment and Baseline Correlation with Treatment Resistance in Metastatic Settings

The pigmentation signature also exhibited dynamic changes and predictive value in the context of ICI treatment and resistance in metastatic melanoma. Analysis of scRNA-seq and bulk RNA-seq data of Pt1 revealed a significant decrease in the pigmentation signature following treatment with aPD1 (**Figure 7**). This decline suggests that ICI therapy may drive or select for dedifferentiation in tumor cells, potentially contributing to the emergence of more aggressive tumor states in certain patients. Among the top differentially expressed genes distinguishing progressors from non-progressors, STXBP6 was identified as a potential mediator of ICI resistance (**Figure 7C**). This gene has been described as an inhibitor of IRF1, a regulator of PD-L1 expression (**Figure 7E**)^26^. Consistent with these findings, our bulk RNA-seq data demonstrated a negative correlation between STXBP6 and PD-L1 expression (**Figure 7D**) further implicating STXBP6 in modulating immune escape mechanisms. In our ITM cohort (stage III disease), high pigmentation signature scores were associated with a reduced risk of distant progression, supporting their role as a favorable prognostic marker. Importantly, while we did not have sufficient numbers to assess the relationship between pigmentation and treatment response in our ITM cohort, analysis of an independent cohort of metastatic melanoma patients (stage IV) treated with anti-PD1 therapy revealed that high pigmentation signature scores were significantly associated with treatment resistance (**Figure 7F**). This contrast suggests that pigmentation reflects a differentiated tumor state that is linked to improved outcomes in earlier-stage disease but may also mark a subset of tumors that are intrinsically less responsive to immune checkpoint blockade in the metastatic setting, highlighting its context-dependent clinical implications.

**Figure 7.**
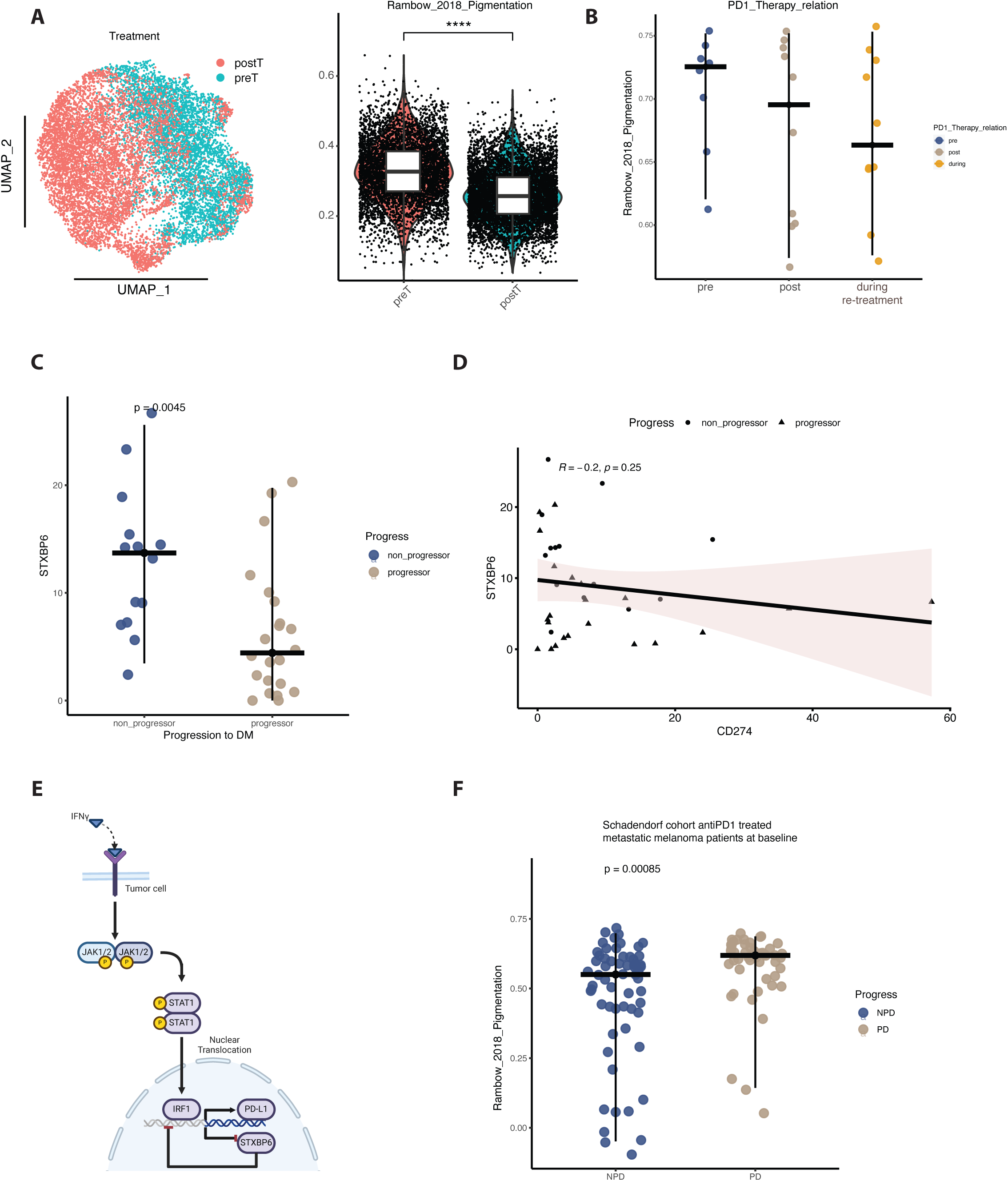
Pigmentation Signature Dynamics and Treatment Resistance in Melanoma. **A.** Pigmentation signature analysis using scRNAseq data from Patient 1 (Pt1), comparing pre-treatment and post-treatment samples. **B.** Box plot comparing pigmentation signature values across pre, post, and during anti-PD1 treatment timepoints in Pt1. **C.** Comparison of STXBP6 expression between non-progressors and progressors in the cohort. **D.** Correlation plot showing the relationship between STXBP6 expression and CD274 (PD-L1) levels across samples in the cohort. **E.** Schematic representation of the suggested functional role of STXBP6 in treatment resistance pathways. **F.** Rambow pigmentation signature comparison between progressors and non-progressors in a publicly available cohort of metastatic melanoma patients treated with anti-PD1 therapy.

## DISCUSSION

In this study, we identify enrichment of melanoma dedifferentiation and T cell exhaustion signatures of melanoma with in-transit metastases progressing to distant metastasis. We leveraged multi-omic profiling of longitudinal samples from two ITM patients, alongside a cohort of 41 additional baseline ITM cases, to identify a pigmentation related signature as a key feature for favorable outcome. Moreover, we determine patient-specific genomic trajectories leading to contrasting clinical outcomes, and provide insights into the impact of PD-1 blockade in ITM. Melanoma tumor dedifferentiation has been associated with resistance to both targeted-and immuno-therapies and worse prognosis ^20,27–30^, but these studies have primarily focused in late stage disease where tumor dedifferentiation is more commonly detected. To our knowledge, our study is the first to demonstrate that markers of dedifferentiated tumor states at earlier stages of melanoma predict progression to distant disease. This addresses a critical need for prognostic biomarkers at earlier stages of disease, where improved prognostic tools would help identify which patients would differentially benefit from often toxic systemic therapy in addition to curative-intent local therapy.

ITM represent a distinct pattern of melanoma spread, occurring between the site of the primary tumor and the first regional lymph node basin. Clinically, ITM can present in various forms, ranging from numerous small, uncountable skin and subcutaneous lesions to large, unresectable tumor masses. Approximately 20% of patients with cutaneous melanoma develop some form of ITM, and treatment options depend on both the stage and tumor burden. This study specifically focuses on unresectable ITM cases in whom surgical intervention could not render the patient free of disease. Molecular features of ITM have not previously been extensively studied, and here, we provide data for future biomarker testing, to identify patients at the highest risk of developing distant metastases.

The clinical data in our study was well balanced: 22 out of 41 patients progressed to distant stage IV disease, while 19 remained in unresectable stage III (non-progressors). Non-progressors were significantly older than those who progressed, whereas tumor characteristics—such as histological subtype and Breslow thickness—were relatively well balanced across both groups. Most patients had primary tumors located on the lower extremities, with a notably high proportion (20%) of the acral lentiginous melanoma (ALM) subtype and lower extremity primaries (54%), consistent with previous reports on ITM development and underlining the importance of biological context in this patient subgroup^2,5^. Pt1 in our study with an ALM primary already presented multiple features of aggressive growth within the primary melanoma (clefting, ulceration, and poorly differentiated morphology), however a high morphological heterogeneity within the primary and among the longitudinal samples was present, which in combination with a minor immune response could explain the patients’ aggressive disease course, but not necessarily why ITM diverge in the first place.

Although the local response to anti-PD1 treatment in our cohort of ITM patients appears limited as evidenced by stable or progressive disease in the majority of treated cases, our data suggest that immune checkpoint blockade may still play a critical role in preventing progression to stage IV disease. In our cohort, first line treatment of unresectable ITM with anti-PD1 was significantly associated with non-progression to distant metastasis, but local response to therapy was not necessary for durable non-distant progressive disease. This aligns with the concept of a disconnected local and systemic immune response, where the tumor microenvironment remains “immune cold” or immunosuppressed, but systemic immune activation is sufficient to prevent dissemination or control occult micro-metastases. These observations mirror the improved distant metastasis-free survival (DMFS) reported in adjuvant anti-PD1 trials for stage III melanoma (Keynote-054, CheckMate-23824,25), which are numerically higher, highlighting the importance of timing and context in immunotherapy efficacy.

The timing of immunotherapy relative to tumor evolution may be especially critical. In Pt1, NGFR+ and AXL+ cells were already present in the primary tumor and seven tumor lineages were already present at the time of anti-PD1 initiation, suggesting that mechanisms of immune escape and clonal complexity were well established when ICI was started. Furthermore, the brain metastasis in Pt1 diverged early in molecular evolution but emerged late in the disease course, and was characterized by a clonal oncogenic NF1 stop codon mutation, while Lineage 7 (Pt1-L7) carried an ARID2 loss-of-function mutation, reflecting a potential role in immune evasion or therapy resistance^31^, underlining how heterogeneity within the primary melanoma shapes disease course and pattern of disease at an early point in time. In contrast, Pt2 developed most lineages during and as results of chemotherapy, which may have induced a highly immunogenic tumor environment, as reflected by increased TMB and TCR diversity, ultimately contributing to complete remission without subsequent systemic therapy. These cases highlight how clonal evolution, therapeutic timing, and immune contexture intersect to influence outcome. From a clinical standpoint, early systemic therapy initiation goes in line with the ongoing paradigm shift towards treating patients systemically with neoadjuvant therapy before tumor surgery is performed which has shown groundbreaking results for stage III disease and is currently trialed during primary diagnosis (^32,33^ NCT05418972 and MARIANE trial NCT06240143). Moreover, the importance of the genomic characteristics of the primary melanoma in predicting recurrence and informing adjuvant anti-PD1-based ICI use is currently being investigated in the NivoMela trial (NCT04309409), whose data may enable direct testing of this hypothesis.

Mechanistically, several factors may underlie the limited local response to PD-1 therapy in ITM. Acral melanomas, such as in Pt1, show lower response rates and worse outcomes compared to cutaneous melanoma and are often associated with reduced angiogenesis and perfusion, which could impair immune cell infiltration and drug delivery in cutaneous tissues^34–36^. Additionally, recent studies have pointed to physical barriers such as cancer-associated fibroblasts (CAFs), which may localize at tumor margins and block T cell entry, particularly in patients with innate resistance to immunotherapy^12^. Furthermore, spatial transcriptomic analyses have shown significant regional heterogeneity within tumors, even within the same lymph node metastasis^37^, reinforcing the need for spatially resolved profiling in understanding treatment resistance. In our recent work^38^, we demonstrated that acral melanomas possess significantly higher genomic heterogeneity, which was positively correlated with COSMIC Signature 3 (indicative of defective homologous recombination repair) and negatively with the UV-associated COSMIC Signature 7, providing a rationale for the high genomic complexity observed in non–sun-exposed tumors. These findings suggest a convergence of genomic instability, proliferative potential, and immune evasion in driving the aggressive clinical course of acral melanoma.

Chemotherapy played a major role in reshaping the immune environment in Pt2. Indeed, dacarbazine exposure induced a mutational signature associated with alkylating agents and resulted in an increase in TMB and TCR diversity, hallmarks of an enhanced neoantigen landscape and immune priming in Pt2. This supports the hypothesis that chemotherapy in ITM patients may not only debulk tumors but in some cases may also facilitate an effective immune response, particularly in combination or sequence with immunotherapies^39^. Data from several randomized phase II and III trials as well as real-life datasets indicate that chemotherapy-based treatments are less effective than ICI-based therapies for advanced melanoma. As a result, chemotherapy is now considered a secondary option, typically reserved for third-line or later treatments, if used at all^40,41^.

Despite the strength of longitudinal and spatial profiling in our study, we acknowledge limitations in statistical power due to sample size. Larger cohorts are needed to validate the prognostic value of the pigmentation and exhaustion signatures and to better characterize systemic versus local immune effects of immunotherapy. From a translational perspective, our findings suggest that pigmentation scoring on standard H&E slides, along with MART1, MKI67 and PCNA marker profiling, could serve as practical tools to stratify patients at risk of progression. These markers correlated closely with the transcriptomic pigmentation signature and immune infiltration (**Supplementary Figure 10**), offering a feasible path toward routine pathology-based prognostic assessment in ITM melanoma.

In conclusion, our study highlights the complex interplay between tumor differentiation, immune exhaustion, and systemic immune dynamics in ITM melanoma. These insights underscore the need for personalized approaches that consider tumor evolution, spatial context, and immune status when selecting and timing therapies for patients with advanced stage III melanoma. The differences observed in this study regarding the treatment response of local versus distant metastases should be the focus of further investigations, preferably prospective in design.

## ACKNOWLEDGEMENTS

A.Z. was supported by the BMBF in the framework of the BMBF Advanced Clinician Scientist Programme UMEA²; 01EO2104. D.L. is funded by the Melanoma Research Alliance (Young Investigator Award). F.R. is funded by the Melanoma Research Alliance (Young Investigator Award) and the Wolfgang & Gertrud Boettcher Foundation. Part of this work was funded by the Deutsche Forschungsgemeinschaft (DFG, German Research Foundation) - SCHA 422/17-1. G. T. was supported by an American-Italian Cancer Foundation Post-Doctoral Research Fellowship. **DATA AND**

## SOFTWARE AVAILABILITY

All analyzed data are in supplementary tables or data available on github at https://github.com/davidliu-lab

## METHODS

### Patient cohort and clinical end points

Patients were identified in databases of participating sites. For enrollment, patients were required to have unresectable melanoma due to in-transit disease (+/− lymph node involvement) without distant metastasis at diagnosis of unresectable melanoma disease. Tissue obtained from in-transit metastasis was required for enrollment and was collected during routine medical care. Clinicopathological and demographic data were collected from patient records locally and are shown in Table 1. Systemic therapy and first line anti-PD1-based therapy were documented for unresectable stage III due to in-transit disease. BOR to anti-PD1-based therapy was assessed according to RECIST criteria v.1.1 by the participating sites. Patients without progression to stage IV (distant metastasis) were grouped as non-progressors, whereas patients showing PD to distant sites were referred to as progressors. Patients were classified as MR when achieving unequivocal responses in individual existing lesions but also progression in others or new lesions. OS was defined as the time between the diagnosis of stage III unresectable and the date of death (any cause). For subjects without documentation of death, OS was censored on the last date the patient was known to be alive. This retrospective study and associated informed consent procedures were approved by the central Ethics Committee (EC) of the University Hospital Essen (18-8123-BO, 12-5152-BO and 11-4715). Approval by the local EC was obtained by investigators if required by local regulations. The list of all the ITM samples at baseline with the clinical annotation is presented in Supplementary Table 1.

### Clinical history and sample collection

Samples were collected retrospectively and obtained by excision or biopsy of melanoma tissue, collected locally at the participating sites and provided formalinfixed and paraffin-embedded (FFPE). Samples were resected between March 2007 and December 2018. All biopsies from the cohort were from in-transit metastasis. Biopsies from the longitudinal single patients approach are as described in the manuscript.

#### Patient 1

A 59-year-old female was diagnosed with stage IIIB acral lentiginous melanoma located on the left palm. Initial treatment included wide local excision and sentinel lymph node biopsy, which revealed metastasis in a left inguinal lymph node. One year later, the patient developed a recurrence with a left inguinal lymph node metastasis. She declined adjuvant interferon-gamma therapy. Imaging showed no further metastases, and a lymphadenectomy was performed, revealing a 2.2 cm lymph node metastasis without evidence of extracapsular extension.

Four months post-lymphadenectomy, the patient developed the first in-transit metastasis (ITM) in the left plantar region, which was surgically excised. Over the subsequent six months, multiple additional ITMs emerged on the left foot, lower leg, and thigh, rendering the disease unresectable. PET-CT imaging revealed bilateral pulmonary nodules and confirmed the presence of multiple in-transit and satellite metastases on the plantar foot and left leg.

The patient was enrolled in a phase Ib/II clinical trial investigating LEE011 (CDK4/6 inhibitor) and MEK162 (MEK inhibitor) for patients with NRAS-mutant melanoma. However, treatment was discontinued after four weeks due to patient preference in the context of grade 1 toxicity. Multiple resections of ITMs were performed both prior to and following systemic therapy.

The patient opted against further systemic therapy and declined isolated limb perfusion. Instead, she underwent multiple additional surgical resections at her own request. Twelve months later, progression in the left iliac/inguinal region required further surgical intervention, which resulted in an R1 resection margin. Anti-PD-1 therapy with pembrolizumab was resumed. Follow-up imaging after four treatment cycles demonstrated progression in both the lungs and left leg skin. Therapy escalation was advised but declined; pembrolizumab monotherapy was continued. Six months into treatment, imaging revealed cerebellar metastases (stage M1d) along with further systemic disease progression.

Cerebral surgery was recommended for symptomatic brain metastases, and therapeutic intensification was again discussed. Nevertheless, the patient chose to continue anti-PD-1 therapy beyond progression and declined brain radiotherapy. Additional surgical procedures were carried out to manage locoregional cutaneous progression on the left leg and lower abdomen.

Subsequent imaging revealed gastric wall thickening, and further diagnostic work-up confirmed the presence of a gastric melanoma metastasis, alongside newly developed ITMs on the left leg. Clinical trial enrollment was offered but declined by the patient, who continued pembrolizumab therapy despite evidence of oligo-progression (bipulmonary, gastric, and distant cutaneous metastases).

With continued disease progression, therapy was escalated to combined immune checkpoint blockade with ipilimumab and nivolumab. After one cycle, treatment was de-escalated to nivolumab monotherapy due to patient preference. However, disease continued to progress. Systemic therapy was ultimately discontinued, and the patient transitioned to best supportive care. The patient died seven years after the initial melanoma diagnosis due to progressive metastatic melanoma. The list of all the samples together with the clinical annotation is presented in Supplementary Table 2.

#### Patient 2

A 67-year-old female was initially diagnosed with stage IIB cutaneous melanoma located on the right knee. Two years following initial treatment, the patient developed a local recurrence, which was managed with wide local excision. Six months later, multiple cutaneous satellite metastases appeared, and a biopsy of the left inguinal lymph nodes confirmed metastatic melanoma.

The disease showed locoregional progression with further cutaneous metastases, necessitating multiple surgical excisions. Systemic therapy was initiated as part of a clinical trial with dacarbazine and PTNZK, followed by dacarbazine monotherapy. Despite systemic therapy, the patient continued to develop cutaneous and subcutaneous metastases on the right upper and lower leg, which were managed surgically.

Due to ongoing progression, isolated limb perfusion (ILP) of the right leg was performed using tumor necrosis factor-alpha (TNF-α) and melphalan. This was followed by surgical excision of residual locoregional disease, including bilateral inguinal lymph node and subcutaneous metastases.

Over the subsequent six years, the patient underwent multiple local excisions for recurrent in-transit metastases. The final in-transit metastasis was surgically removed 11 years after the initial melanoma diagnosis. Thereafter, the patient remained disease-free, with no evidence of local or distant recurrence during a nine-year period of clinical and radiological follow-up. The list of all the samples together with the clinical annotation is presented in Supplementary Table 3.

### Whole-exome and whole-transcriptome sequencing

DNA extraction, whole exome library preparation and sequencing were performed for samples as previously described (^42–44^). Slides were cut from FFPE blocks and macrodissected for tumor-enriched tissue. Paraffin was removed from FFPE sections and cores using CitriSolv (Fisher Scientific), followed by ethanol washes and tissue lysis overnight at 56 °C. Samples were then incubated at 90 °C to remove DNA cross links. Extraction of DNA, and, when possible, RNA was performed using the QIAGEN AllPrep DNA/RNA mini kit (51306). Germline DNA was obtained from peripheral blood mononuclear cells and adjacent normal tissue. Whole-exome sequencing libraries were prepared using Twist library preparation kits and captured using the Twist exome panel, then sequenced on an Illumina platform with paired-end 150 bp reads. Quality control and somatic variant calling were performed using the CGA WES Characterization Pipeline developed by the Getz Lab at the Broad Institute (https://github.com/broadinstitute/CGA_Production_Analysis_Pipeline) as previously described^38,42,43^.

Bulk RNA-sequencing was performed for the baseline ITM tumor samples. FASTQ files were assessed using FASTQC (v0.11.9; http://www.bioinformatics.babraham.ac.uk/projects/fastqc/) to evaluate quality metrics. STAR (v2.7.0)^45^ was used for alignment to the hg19 reference genome, and quantification was done using Salmon (v0.14.1)^46^. TPM values were normalized across samples, and only protein-coding genes expressed in at least 5% of samples were retained. Samples with fewer than 1 million reads or <50% uniquely mapped reads were excluded. Additional outliers were identified by principal component analysis (PCA) of alignment and expression metrics. Three samples were discarded due to multiple failures in the QC () with a total of 36 samples passing the QC.

BulkRNAseq downstream analysis was implemented in R using the package GSVA (71)(v 1.44.0) and msigdbr (72)(v7.5.1). Tumor and Immune states evaluated were obtained from recently published works^19,23^. Bulk deconvolution and tumor compartment identification were performed with bayesprism (https://www.bayesprism.org/).

### Phylogenetic analysis

Somatic variant calls, allelic copy number data, and sample purity/ploidy estimates were used to infer clonal structures and phylogenies using PyCloneVI^47^ and PhylogicNDT^48^. For each patient, merged MAF files were generated, and force-calling was performed across all samples to assess variant presence. ABSOLUTE^49^ was used to estimate cancer cell fractions (CCFs) for each mutation. PyCloneVI, a Bayesian clustering algorithm, was applied to assign mutations to clusters based on their CCFs and copy number context. For Pt1, 14 clusters were identified, with 9 retained after filtering for clusters containing >3 mutations. For Pt2, 15 clusters were initially identified, and 11 retained after filtering. PhylogicNDT was then used to build patient-specific phylogenetic trees and evolutionary trajectories, integrating clonal structure and temporal data.

#### Patient1

Cluster 10 contained the NRAS mutation along with other clonal alterations and was detected in all tumor samples, forming the clonal trunk of the phylogeny. This cluster defined Lineage 1, which included the primary tumor and regional lymph node samples and represented the earliest detectable ancestral population.

A split into two evolutionary trajectories was then observed, separating early-stage from late-stage disease. Cluster 1 mutations, present only in early-stage samples, defined the early evolutionary branch. Cluster 9 mutations were exclusive to the late-stage samples, marking the divergence of more advanced disease.

Among early-stage lineages, four were defined:

- Lineage 1: Contained only clonal mutations from cluster 10.
- Lineage 2: Characterized by cluster 6, enriched exclusively in the brain metastasis sample (T22), which had the highest number of newly acquired clonal mutations.
- Lineage 3: Defined by mutations in cluster 12, observed only in ITM samples.
- Lineage 4: Defined by mutations in cluster 7, also confined to ITM samples.

Among late-stage lineages, three were identified:

- Lineage 5: Included samples harboring both clonal mutations (cluster 10) and late-stage associated mutations (cluster 9), but no additional private alterations.
- Lineage 6: Built upon Lineage 5 with additional mutations from cluster 11.
- Lineage 7: Derived from Lineage 5 and enriched for alterations in cluster 4.

This phylogenetic structure supports a model in which early and late-stage metastatic disease evolved from a common ancestor but accumulated distinct alterations that shaped their clonal trajectories, with Lineage 5 as the only lineage appearing before and after treatment and the only lineage constituted by both ITM and distant metastasis samples.

#### Patient2

For Patient 2, the primary tumor sample was not available; the earliest sample analyzed was an in-transit metastasis (ITM). PyCloneVI identified that mutations from clusters 2 and 3, including a clonal BRAF V600E mutation in cluster 2, were shared across all samples, forming the trunk of the phylogeny. Subsequent branching was driven by alterations in cluster 13, defining Lineage 2, and in cluster 4, defining Lineage 1. Early samples (T1, T2, T7), obtained before chemotherapy, clustered with Lineage 1 and were enriched for the UV mutational signature. This lineage also included samples obtained during chemotherapy (T9, T10), where a marked shift occurred: the appearance of COSMIC Signature 11 associated with alkylating agents led to a drastic increase in tumor mutational burden.

From Lineage 1, several additional branches emerged due to the increased mutagenesis:

- Lineage 5: Defined by mutations in cluster 12.
- Lineage 6: Characterized by alterations in cluster 1.
- Lineage 7: Characterized by alterations in cluster 6.

In parallel, from Lineage 2, additional evolution produced:

- Lineage 3: Defined by mutations in cluster 14.
- Lineage 4: Defined by alterations in cluster 10.

These findings suggest that chemotherapy-induced mutagenesis contributed to the emergence of several evolutionary branches in Patient 2, reflecting the adaptive potential of the tumor under therapeutic pressure.

### Imaging (H&E and tissue-based CyCIF)

H&E-stained FFPE sections from each tissue block were imaged using a CyteFinder slide scanning fluorescence microscope (RareCyte Inc.) with a 20×/0.75 NA objective with no pixel binning. Serial FFPE sections (5 μm thick) were subjected to whole-slide, subcellular-resolution, XX-plex CyCIF imaging. CyCIF was performed as described in Lin et al. 2018 ^13^. In brief, the BOND RX Automated IHC Stainer was used to bake FFPE slides at 60°C for 30 minutes, dewax using Bond Dewax solution at 72°C, and perform antigen retrieval using Epitope Retrieval 1 (Leica) solution at 100°C for 20 minutes. Slides underwent multiple cycles of antibody incubation, imaging, and fluorophore inactivation. Antibodies were incubated overnight at 4°C in the dark. Before imaging, a 0.15mm single-sided self-adhesive spacer was applied to the slide, followed by wet-mounting of a glass coverslip using 50% glycerol in 1× PBS. Images were acquired using the same microscope and objective as the H&E images. Slides were soaked in 42°C PBS to facilitate coverslip removal, and then fluorophores were inactivated by incubating slides in a solution of 4.5% H_2_O_2_ and 24 mmol/L NaOH in PBS and placing them under an LED light source for 1 hour. The list of all antibody panels used in the experiments is presented in Supplementary Table 4-7.

### CyCIF image pre-processing and quality control

CyCIF image pre-processing (stitching, single cell segmentation, feature extraction) was performed using the MCMICRO pipeline^50^. Multiple approaches were also taken to ensure the quality of the single-cell data. At the image level, the cross-cycle image registration and tissue integrity were reviewed; regions that were poorly registered or contained severely deformed tissues and artifacts were identified, and cells inside those regions were excluded.

### Single-cell spatial analysis

We used a gating-based phenotyping approach to classify cells as described previously in Nirmal et al 2022. In short, an open-source visual gating tool (https://github.com/labsyspharm/gater) was used to derive gates for each marker. After gating, tumor cells were identified using SOX10+ as a marker for tumor. We confirmed this visually, and did not observe tumor regions with a loss of SOX10. Then, we determined tumor cell states by using MART1, AXL and NGFR as key protein markers to indicate distinct states within the dedifferentiation axis.

### GeoMX analysis

GeoMx Digital Spatial Profiling (DSP) data from patient Pt1 sample T15 were processed using *R* to compare spatially distinct tumor regions of interest enriched for either AXL or NGFR expression. Count and segment annotation files were loaded using the *readGeoMx* function from the *standR* package. ROIs were annotated using CyCIF as either AXL-high or NGFR-high. Quality control was performed using *plotRLExpr*, PCA visualization, and per-sample annotations. Data normalization was conducted using the TMM method.

Differential expression analysis between AXL-high and NGFR-high nodules was carried out using the *limma-voom* pipeline. A design matrix was constructed based on group annotations, and contrasts were fitted to test for differentially expressed genes. Genes with an FDR < 0.05 were considered significant. Visualization included volcano plots, and heatmaps of selected melanoma-relevant genes.

### snRNA-seq from cryosections

Nuclei extraction was performed according to the protocol from Jones’ Lab^51^. Tumor material of serveral cryossections was transferred into a glass douncer containing 2 ml of chilled lysis buffer (0.32 M sucrose, 5 mM calcium dichloride, 3 mM magnesium acetate, 2.0 mM EDTA, 0.5 mM EGTA, 10 mM Tris-HCl, pH 8.0, 1 mM DTT and 0.1% Triton X-100. The chemical lysis was reinforced by mechanical homogenization by douncing 10 strokes with pestle A and then 10 strokes with pestle B. The lysate was filtered sequentially through a 100 μm and then 40 μm pluriStrainer®. An extra 3 ml lysis buffer was used to wash the dounce homogenizer and both pestles. The suspension was spun down to remove the lysis buffer (500 g, 5 min, 4C). The nuclei were re-suspended in 5 ml wash buffer (0.32 M sucrose, 5 mM calcium dichloride, 3 mM magnesium acetate, 2.0 mM EDTA, 0.5 mM EGTA, and 10 mM Tris-HCl, pH 8.0) in a 50 ml tube Three washes were performed to remove the debris. After the washes, the nuclear pellet was re-suspended in 1 ml of nuclei storage buffer (0.43 M sucrose, 70 mM potassium chloride, 2 mM magnesium dichloride, 10 mM TrisHCl, pH 7.2 and 5 mM EGTA). Nuclei concentrations were assessed using a LUNA-FL Dual Fluorescence Cell Counter. The nuclei were directly further processed for snRNA-sequencing. Single-cell libraries were generated via the Controller Single-Cell Instrument and the Single Cell 3′ Library & Gel Bead Kit v3.1 and Chip Kit G (10x Genomics) according to the manufacturer’s protocol v3.1Rev D. Briefly, nuclei were suspended in 0.04% BSA–PBS. Nuclei were injected in each channel with a targeted nuclei recovery estimate of 7,000. After generation of nanoliter-scale Gel bead-in-EMulsions (GEMs), GEMs were reverse transcribed in a C1000 Touch Thermal Cycler (Bio Rad) programed at 53°C for 45 min, 85°C for 5 min, and hold at 4°C. After reverse transcription, single-cell droplets were broken and the single-strand cDNA was isolated and cleaned with Cleanup Mix containing DynaBeads (Thermo Fisher Scientific). cDNA was then amplified with a C1000 Touch Thermal Cycler programed at 98°C for 3 min, 12 cycles of (98°C for 15 s, 67°C for 20 s, 72°C for 1 min), 72°C for 1 min, and held at 4°C twice. Subsequently, the amplified cDNA was fragmented, end-repaired, A-tailed and index adaptor ligated, with SPRIselect Reagent Kit (Beckman Coulter) with cleanup in between steps. The post-ligation product was amplified with a C1000 Touch Thermal Cycler programed at 98°C for 45 s, 14 cycles of (98°C for 20 s, 54°C for 30 s, 72°C for 20 s), 72°C for 1 min, and hold at 4°C. The sequencing-ready library was cleaned up with SPRIselect beads. The profiles of the final libraries were checked with the Agilent TapeStation.

### snRNA-seq data analysis

Raw fastq files were processed using CellRanger (version 5.0.1) with the hg19-3.0.0 reference genome for mapping. The resulting gene count matrix was analyzed in R using the Seurat package. Four samples were merged, and high-quality cells were retained based on the following criteria: number of genes (nFeature) between 1000 and 7500, and mitochondrial gene content less than 10%. Potential doublets were identified and removed using DoubletFinder. This process yielded a total of 26,870 cells for further analysis. For data normalization, Seurat’s SCTransform was applied. Batch effects were corrected using Harmony (v1.0.3). Cell clustering was performed using FindNeighbors and FindClusters functions (dimensions = 1:25, resolution = 0.2), resulting in 10 distinct clusters (0-9). The cells were then categorized into malignant cells (clusters 0-3) and microenvironment (TME) cells (clusters 4-9) for separate analyses. Malignant cells were subsetted and underwent further filtering (nFeature > 2000, mitochondrial gene content < 5%), resulting in 12,404 cells. These cells were normalized using SCTransform, with batch effects corrected by Harmony. Dimensionality reduction was performed using RunUMAP (dimensions = 1:20). For TME cells, 3,785 cells were obtained and processed similarly, including normalization, integration, and clustering (dimensions = 1:30, resolution = 0.5), resulting in 15 clusters (0-14). Clusters 2 and 7, exhibiting high expression of melanoma-related genes and high doublet probability scores, were excluded, leaving 3,156 cells. Dimensionality reduction for these cells was also performed using RunUMAP (dimensions = 1:30). Cells were annotated based on common markers for each cell type. For visualization, including violin plots, cell proportion bar plots, and waterfall plots, the SeuratExtend Package (https://github.com/huayc09/SeuratExtend) was utilized. Statistical analyses were conducted using the Wilcoxon test, with p-values adjusted using the Holm method. To explore which ligands in the TME induced phenotypic changes in malignant cells post-treatment, NicheNet was employed to predict top ligands. The expression of these ligands across different cell clusters and their ligand-receptor interactions were analyzed and visualized.

