## Supplementary Figures for "Multiomic profiling of a unique in-transit melanoma cohort identifies melanoma differentiation as predictor of tumor progression and therapy response"

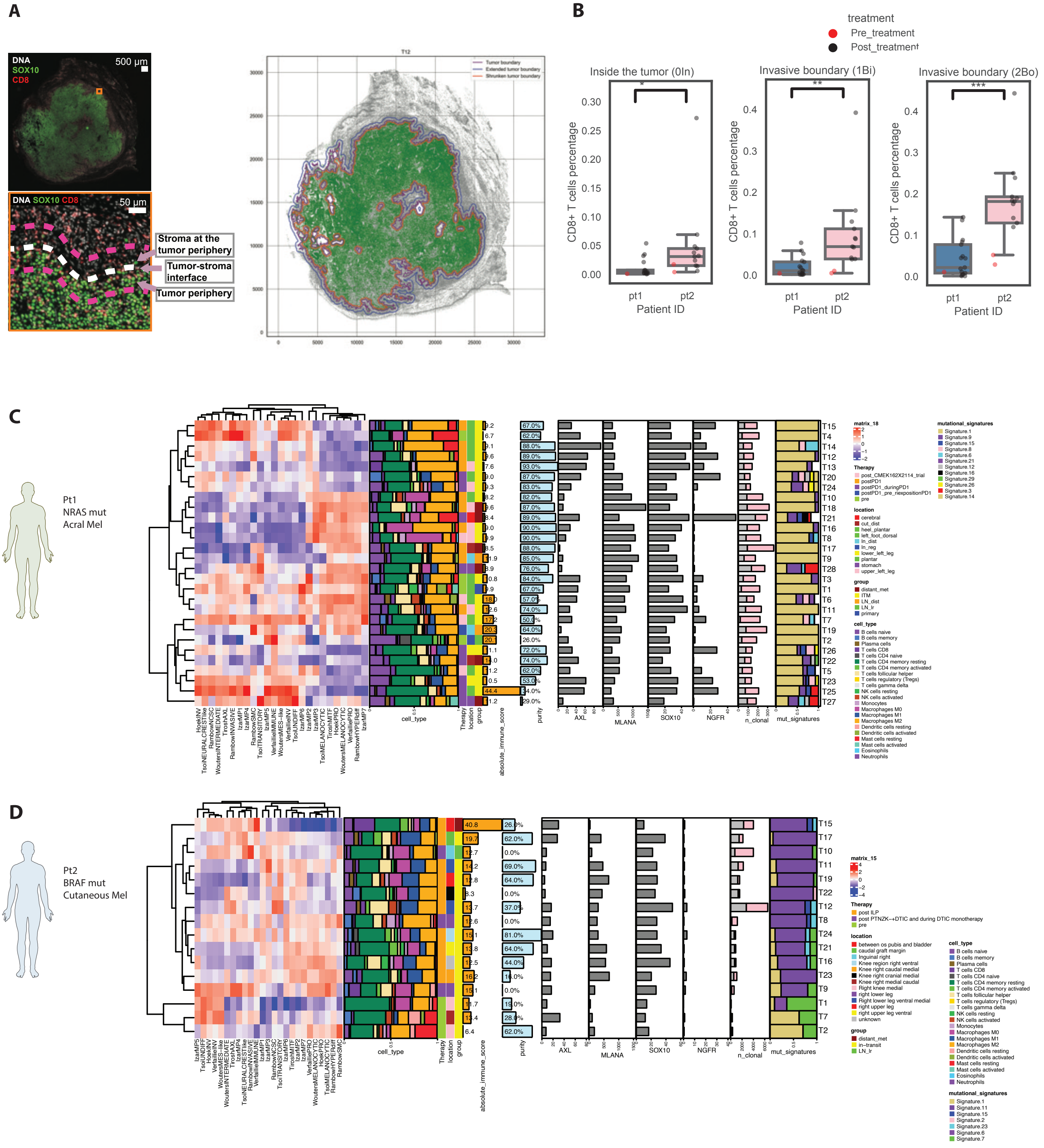
A
DNA
SOX10
CD8
500 µm
DNA SOX10 CD8
50 µm
Stroma at the tumor periphery
Tumor-stroma interface
Tumor periphery
T12
Tumor boundary
Extended tumor boundary
Shrunken tumor boundary
B
treatment
Pre_treatment
Post_treatment
Inside the tumor (0In)
Invasive boundary (1Bi)
Invasive boundary (2Bo)
CD8+ T cells percentage
Patient ID
pt1
pt2
C
Pt1
NRAS mut
Acral Mel
cell_type
Therapy
location
group
absolute_immune_score
purity
AXL
MLANA
SOX10
NGFR
n_clonal
mut_signatures
matrix_18
mutational_signatures
D
Pt2
BRAF mut
Cutaneous Mel
matrix_15

**Supplementary Figure 1. Distinctive Tumor Characteristics of Pt1 and Pt2: Genomic Heterogeneity, CD8+ GZMB+ T Cell Infiltration, Tumor states and mutational signatures.** **A.** CyCIF analysis showing one sample of pt1 and how the tumor boundaries were defined **B.** Comparison of Pt1(in gray) and Pt2 (in pink) T cells CD8+ proportion inside the tumor and in the two invasive boundaries. **C.** Heatmap showing genomic and transcriptomic features of the longitudinal samples of Patient1. **D.** Heatmap showing genomic and transcriptomic features of the longitudinal samples of Patient2.

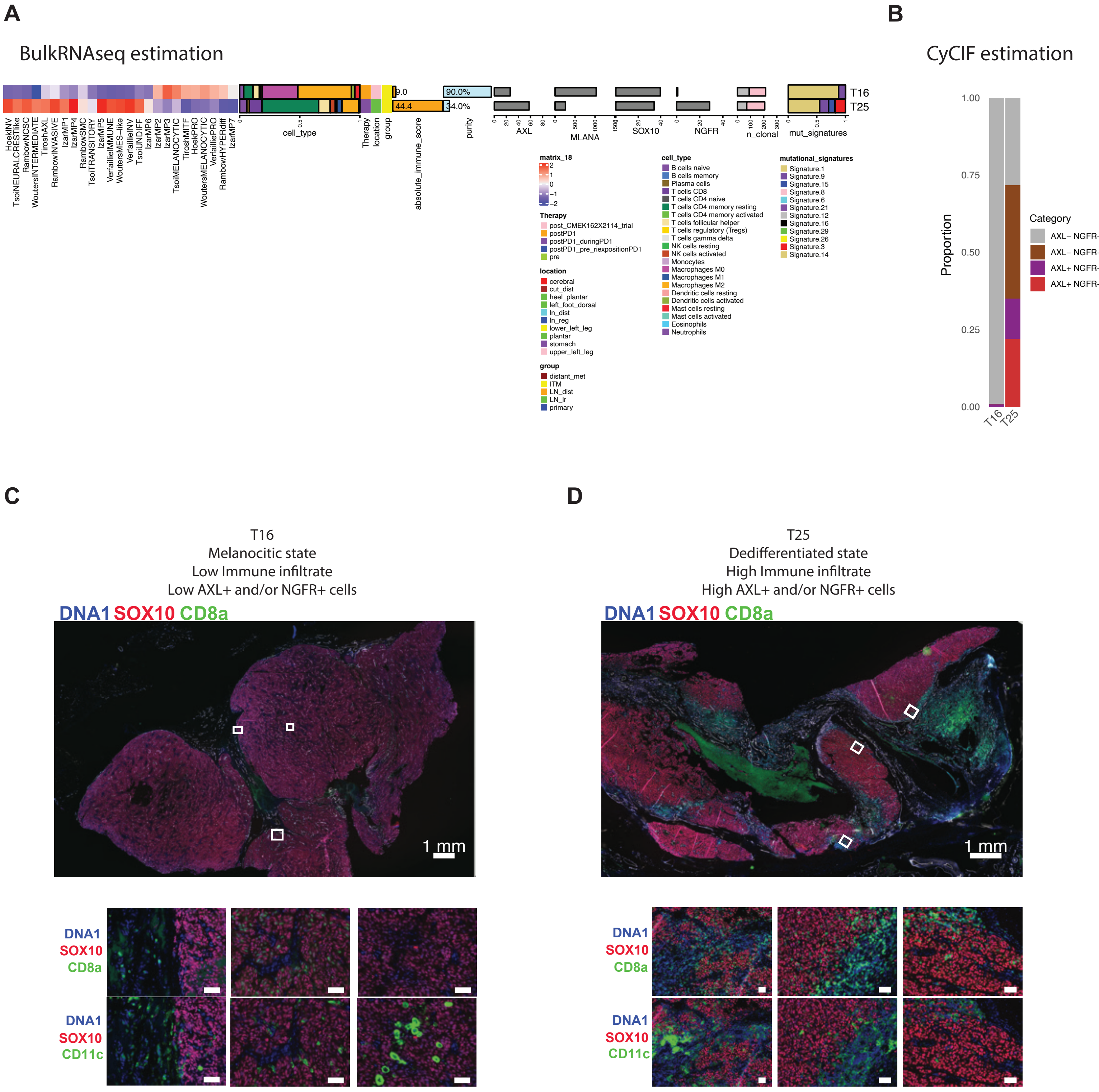
A
BulkRNAseq estimation
T16
T25
cell_type
Therapy
location
group
absolute_immune_score
purity
AXL
MLANA
SOX10
NGFR
n_clonal
mut_signatures
matrix_18
cell_type
mutational_signatures
Therapy
location
group
B
CyCIF estimation
Proportion
Category
AXL− NGFR−
AXL− NGFR+
AXL+ NGFR−
AXL+ NGFR+
C
T16
Melanocitic state
Low Immune infiltrate
Low AXL+ and/or NGFR+ cells
DNA1 SOX10 CD8a
1 mm
DNA1 SOX10 CD11c
D
T25
Dedifferentiated state
High Immune infiltrate
High AXL+ and/or NGFR+ cells
DNA1 SOX10 CD8a
1 mm
DNA1 SOX10 CD11c

**Supplementary Figure 2. The two orthogonal quantifications (bulkRNAseq and CyCIF) showing similar results. A.** BulkRNAseq comparing sample T16 and T25 from patient1, the heatmap include the single sample gene set enrichment analysis, the WES estimation of the purity of the sample, the cibersortx deconvolution of the tumor microenviroment (TME), the cibersort absolute quantification of the immune cells in the TME, the expression AXL, MLANA, SOX10, NGFR the proportion of clonal and subclonal mutation and the mutational signature analysis. **B.** CyCIF estimation of the tumor cells AXL+ and/or NGFR+, double negative cells are SOX10+ (gray). **C.** CyCIF visualization of T16 showing the low immune infiltrate and the high proportion of SOX10+ cells and low/absent AXL+ or NGFR+ cells. **D.** CyCIF visualization of T25 showing the high immune infiltrate and the high proportion of AXL+ or NGFR+ cells.

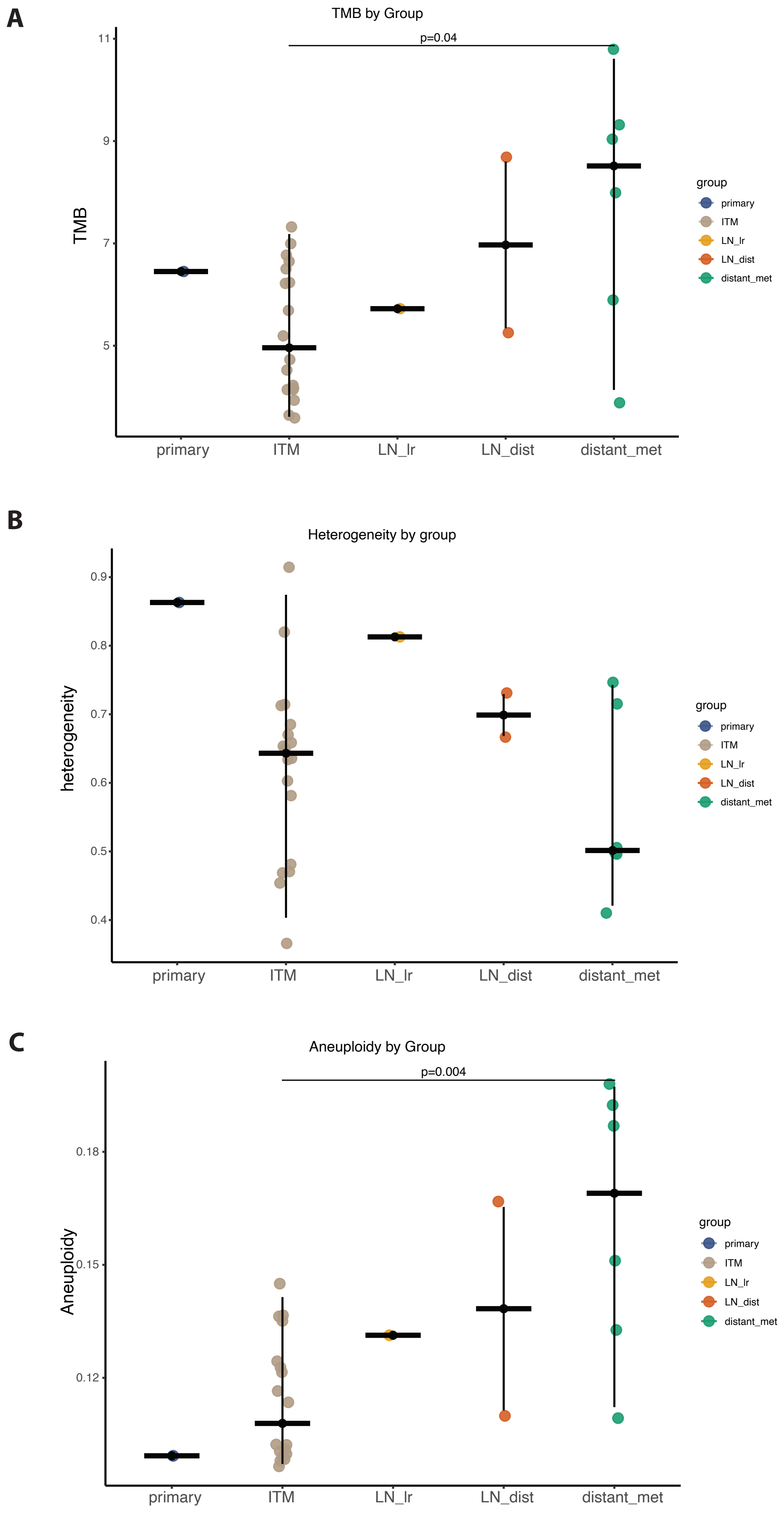
A
TMB by Group
p=0.04
TMB
11
9
7
5
primary
ITM
LN_lr
LN_dist
distant_met
group
primary
ITM
LN_lr
LN_dist
distant_met
B
Heterogeneity by group
heterogeneity
0.9
0.8
0.7
0.6
0.5
0.4
primary
ITM
LN_lr
LN_dist
distant_met
group
primary
ITM
LN_lr
LN_dist
distant_met
C
Aneuploidy by Group
p=0.004
Aneuploidy
0.18
0.15
0.12
primary
ITM
LN_lr
LN_dist
distant_met
group
primary
ITM
LN_lr
LN_dist
distant_met

**Supplementary Figure 3. Comparison between the genomic features of ITM samples and distant metastasis in Pt1. A.** TMB comparison between the different group of samples of pt1, ITM samples showing significant lower tumor mutational burden compared to distant metastasis. **B.** Genomic heterogeneity comparison between the different group of samples of pt1 **C.** Aneuploidy comparison between the different group of samples of pt1, distant metastasis compared to ITM have a significant higher proportion of aneuploidy.

**A**

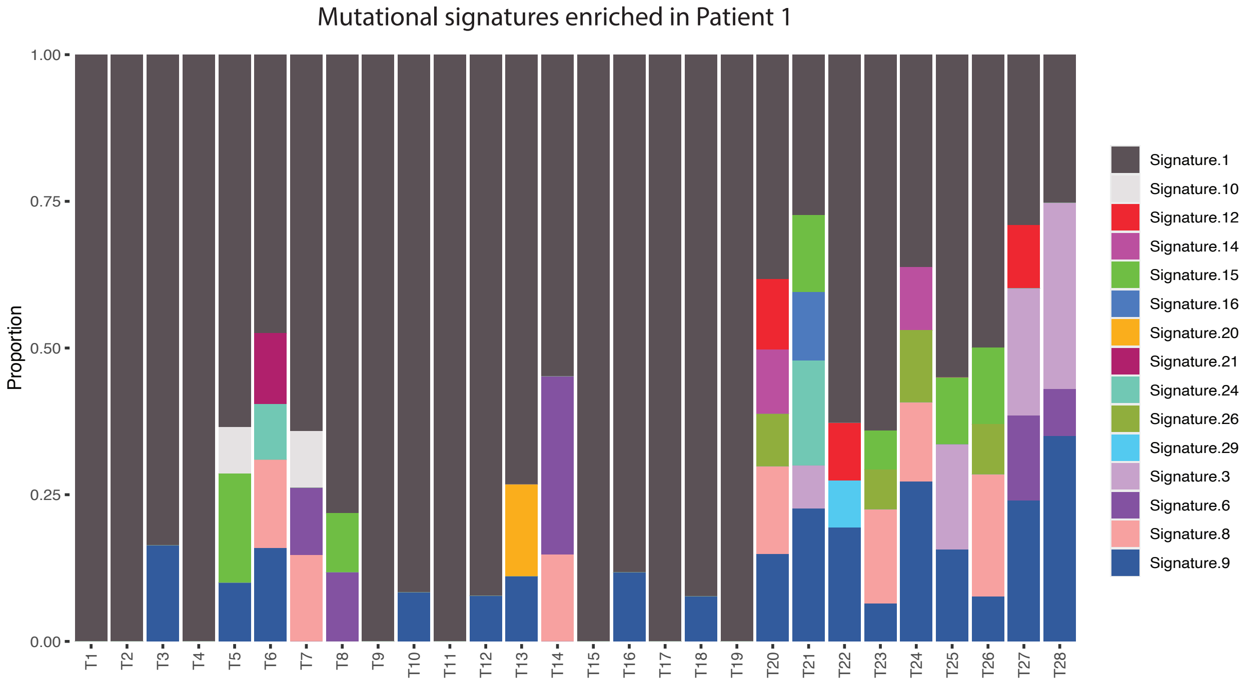
Mutational signatures enriched in Patient 1
Proportion
1.00
0.75
0.50
0.25
0.00
T1 T2 T3 T4 T5 T6 T7 T8 T9 T10 T11 T12 T13 T14 T15 T16 T17 T18 T19 T20 T21 T22 T23 T24 T25 T26 T27 T28
Signature.1
Signature.10
Signature.12
Signature.14
Signature.15
Signature.16
Signature.20
Signature.21
Signature.24
Signature.26
Signature.29
Signature.3
Signature.6
Signature.8
Signature.9

**B**

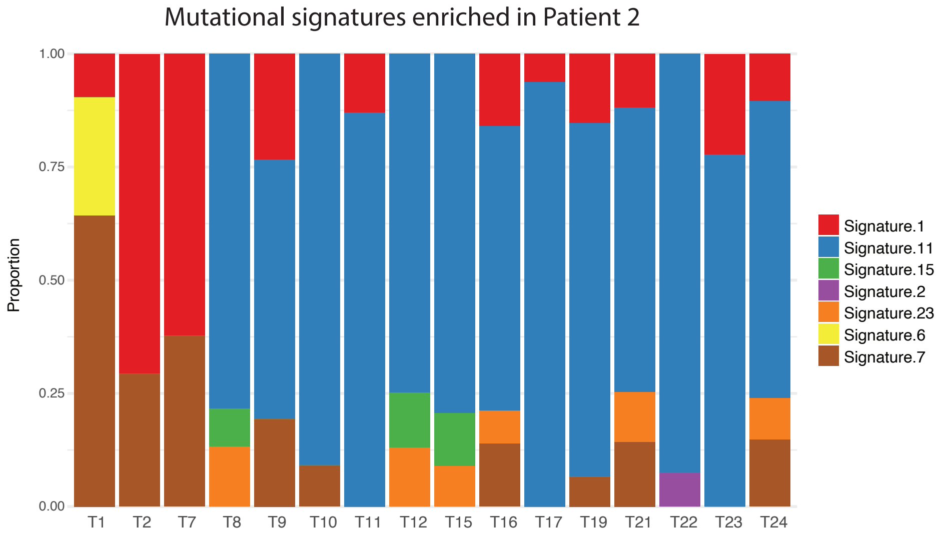
Mutational signatures enriched in Patient 2
Proportion
1.00
0.75
0.50
0.25
0.00
T1 T2 T7 T8 T9 T10 T11 T12 T15 T16 T17 T19 T21 T22 T23 T24
Signature.1
Signature.11
Signature.15
Signature.2
Signature.23
Signature.6
Signature.7

**Supplementary Figure 4. Pt1 and Pt2 COSMIC mutational signature analysis. A.** Pt1 enriched in the aging COSMIC mutational signature**. B.** Pt 2 show enrichment of cosmic signature 11 associated with exposure to alkylating agents.

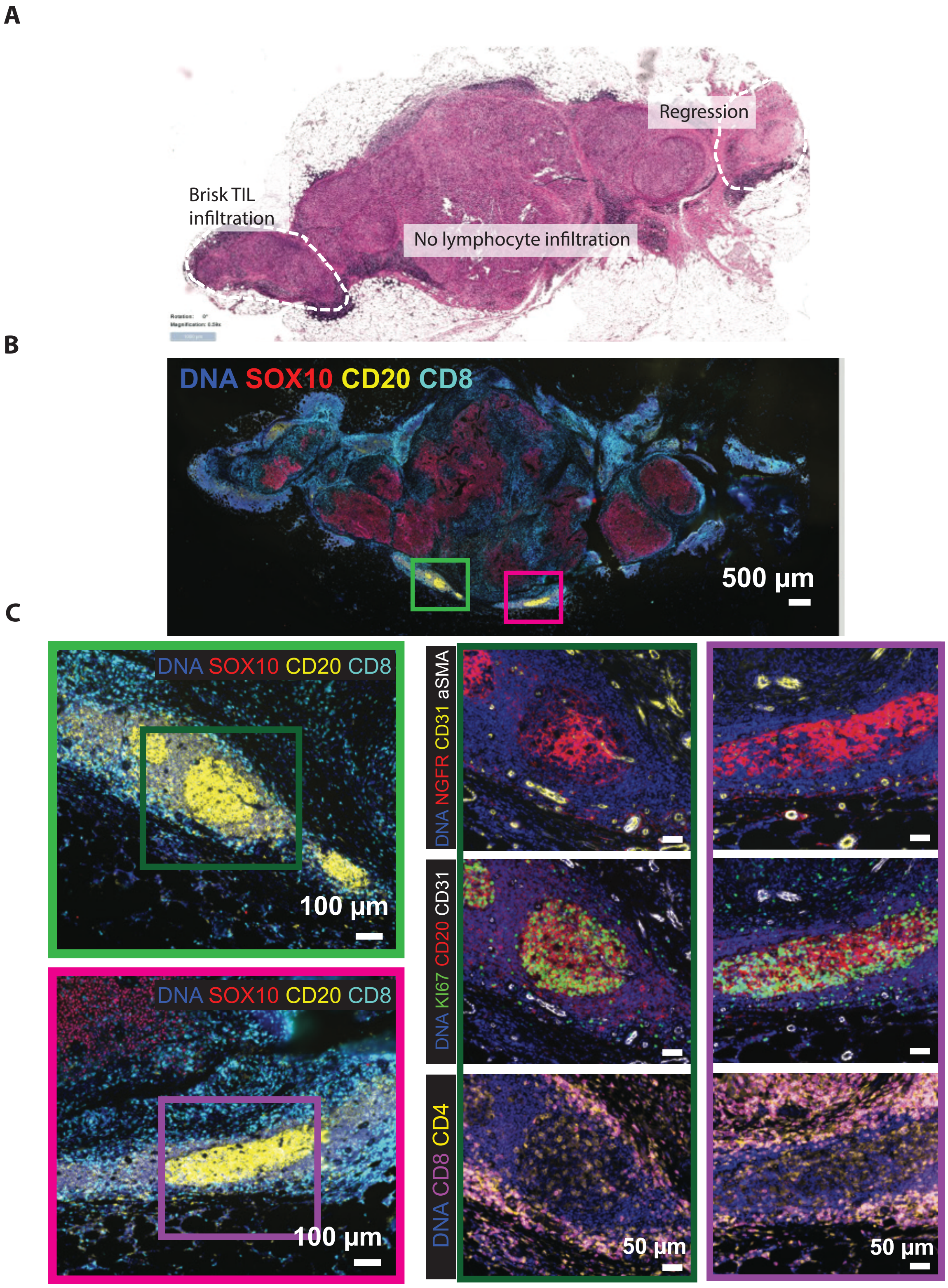

A
Regression
Brisk TIL infiltration
No lymphocyte infiltration
B
DNA SOX10 CD20 CD8
500 µm
C
DNA SOX10 CD20 CD8
100 µm
DNA SOX10 CD20 CD8
100 µm
DNA NGFR CD31 aSMA
DNA KI67 CD20 CD31
DNA CD8 CD4
50 µm
50 µm

**Supplementary Figure 5. Pt2 sample (T25) with features of regression, brisk TIL infiltration and mature tertiary lymphoid structure.**

**A.** H&E staining ofT25 of Pt2. **B.** CyCIF showing T cells (CD8+), B cells (CD20+) and tumor cells (SOX10+), with a high proportion of T cells surrounding the tumor cells. **C.** Two tertiary lymphoid structure with high enrichment of Bcells, Tcells CD8, CD31+ endothelial cells , and high expression of ki67.

A

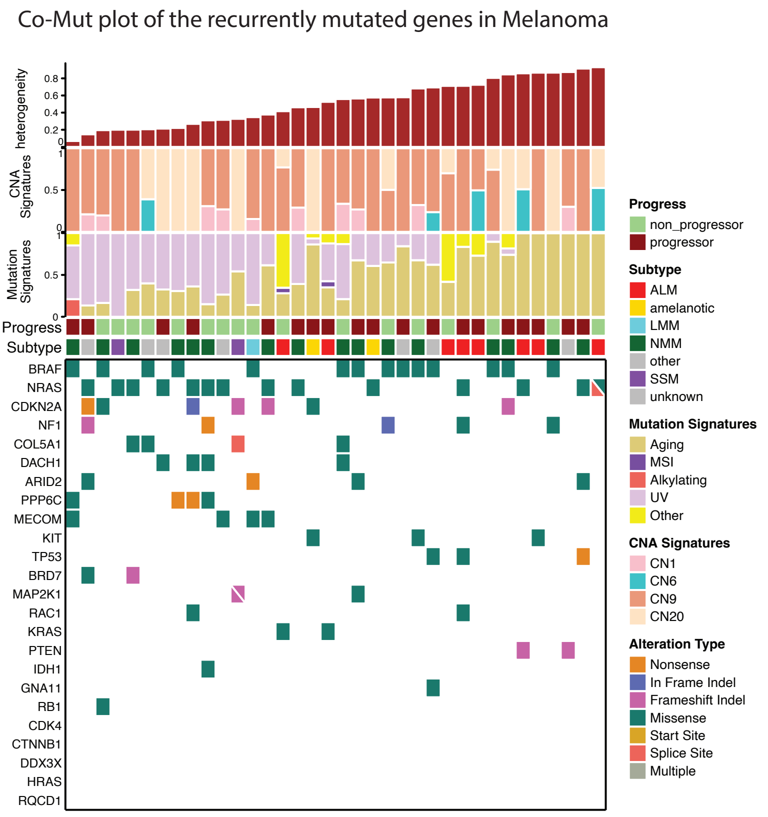
Co-Mut plot of the recurrently mutated genes in Melanoma
heterogeneity
CNA Signatures
Mutation Signatures
Progress
Subtype
BRAF
NRAS
CDKN2A
NF1
COL5A1
DACH1
ARID2
PPP6C
MECOM
KIT
TP53
BRD7
MAP2K1
RAC1
KRAS
PTEN
IDH1
GNA11
RB1
CDK4
CTNNB1
DDX3X
HRAS
RQCD1
Progress
non_progressor
progressor
Subtype
ALM
amelanotic
LMM
NMM
other
SSM
unknown
Mutation Signatures
Aging
MSI
Alkylating
UV
Other
CNA Signatures
CN1
CN6
CN9
CN20
Alteration Type
Nonsense
In Frame Indel
Frameshift Indel
Missense
Start Site
Splice Site
Multiple

B

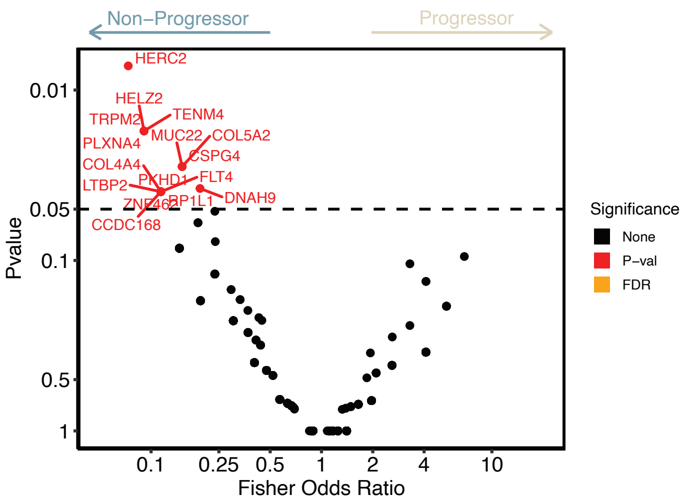
Non–Progressor
Progressor
HERC2
HELZ2
TRPM2
TENM4
PLXNA4
MUC22
COL5A2
CSPG4
COL4A4
PKHD1
FLT4
LTBP2
DNAH9
ZNF462
RP1L1
CCDC168
Pvalue
Fisher Odds Ratio
Significance
None
P–val
FDR

C

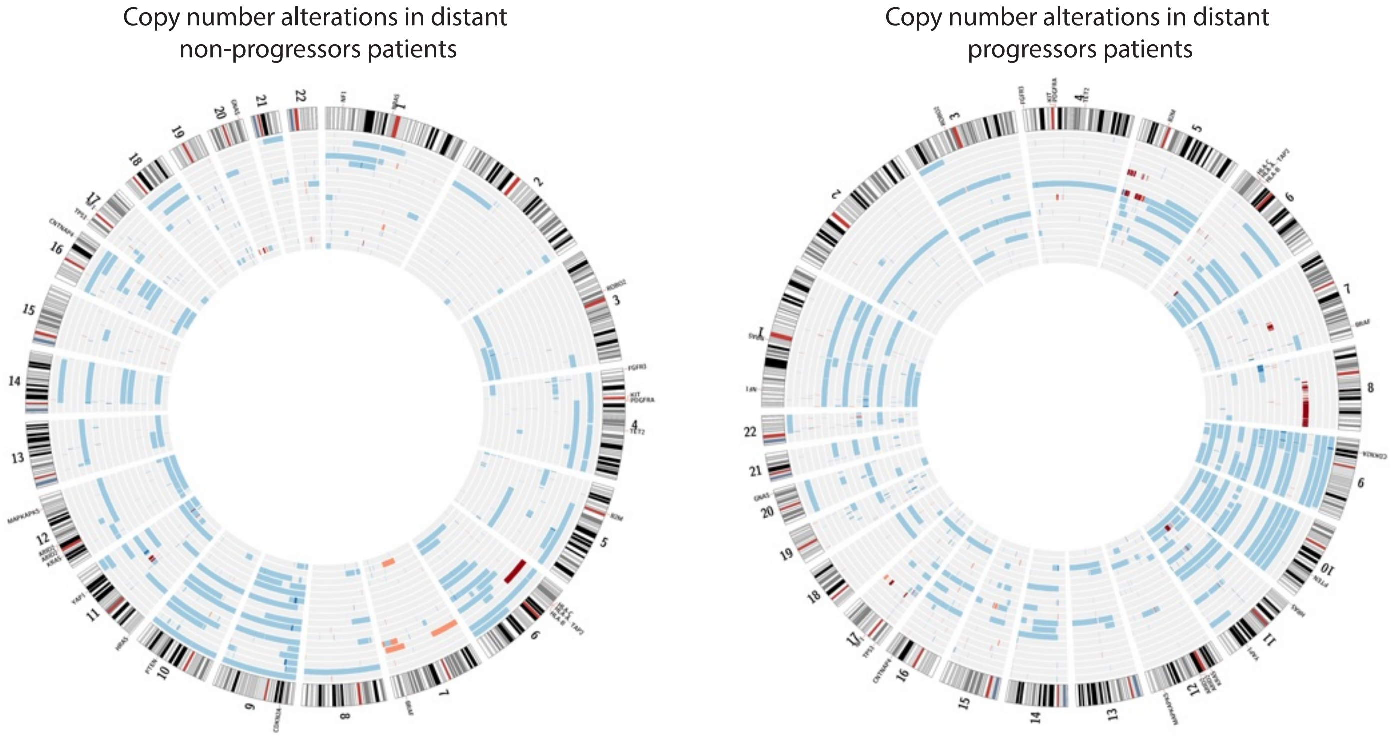
Copy number alterations in distant non-progressors patients
Copy number alterations in distant progressors patients

**Supplementary Figure 6. Cohort genomic characterization. A.** Heatmap showing the commonly mutated genes in Melanoma, together with the proportion of subclonal mutations, mutational signatures analysis and copy number signatures. **B.** Volcano plot showing the mutations enriched in Non-progressors and Progressors, no mutations is significant after multiple hypothesis correction. **C.** Copy number alteration of each patients grouped by non progressors in the left and progressors in the right. Blue indicate homozygous deletion, cyan indicates loss of heterozygosis, light red amplification and dark red high amplification.

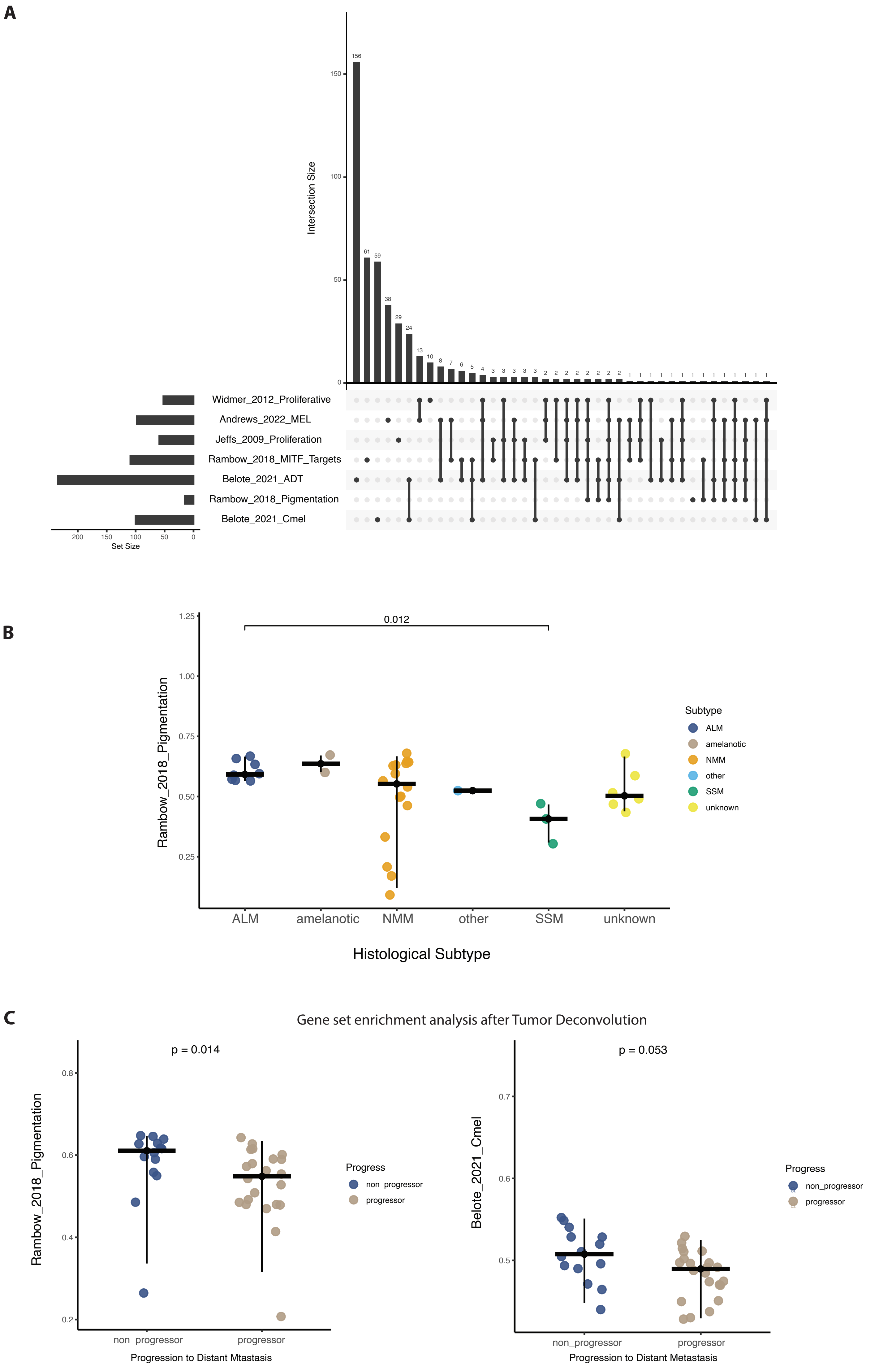
A
Intersection Size
156
61
59
38
29
24
13
10
8
7
6
5
4
Widmer_2012_Proliferative
Andrews_2022_MEL
Jeffs_2009_Proliferation
Rambow_2018_MITF_Targets
Belote_2021_ADT
Rambow_2018_Pigmentation
Belote_2021_Cmel
Set Size
B
0.012
Rambow_2018_Pigmentation
Subtype
ALM
amelanotic
NMM
other
SSM
unknown
Histological Subtype
C
Gene set enrichment analysis after Tumor Deconvolution
p = 0.014
Rambow_2018_Pigmentation
Progress
non_progressor
progressor
Progression to Distant Mtastasis
p = 0.053
Belote_2021_Cmel
Progression to Distant Metastasis

**Supplementary Figure 7. Investigating the Cmel and the Pigmentation signatures.**
**A.** Genes shared between the significant signatures, no genes are shared between the pigmentation and the Cmel signatures. **B.** Boxplot comparing the pigmentation signature enrichment after grouping the patients by Subtype, Acral melanoma posses and higher pigmentation signature. **C.** Single sample gene set enrichment analysis after tumor deconvolution, the Pigmentation signature is even more significant when evaluated only on the tumor component, for the Cmel signature instead there is a trend.

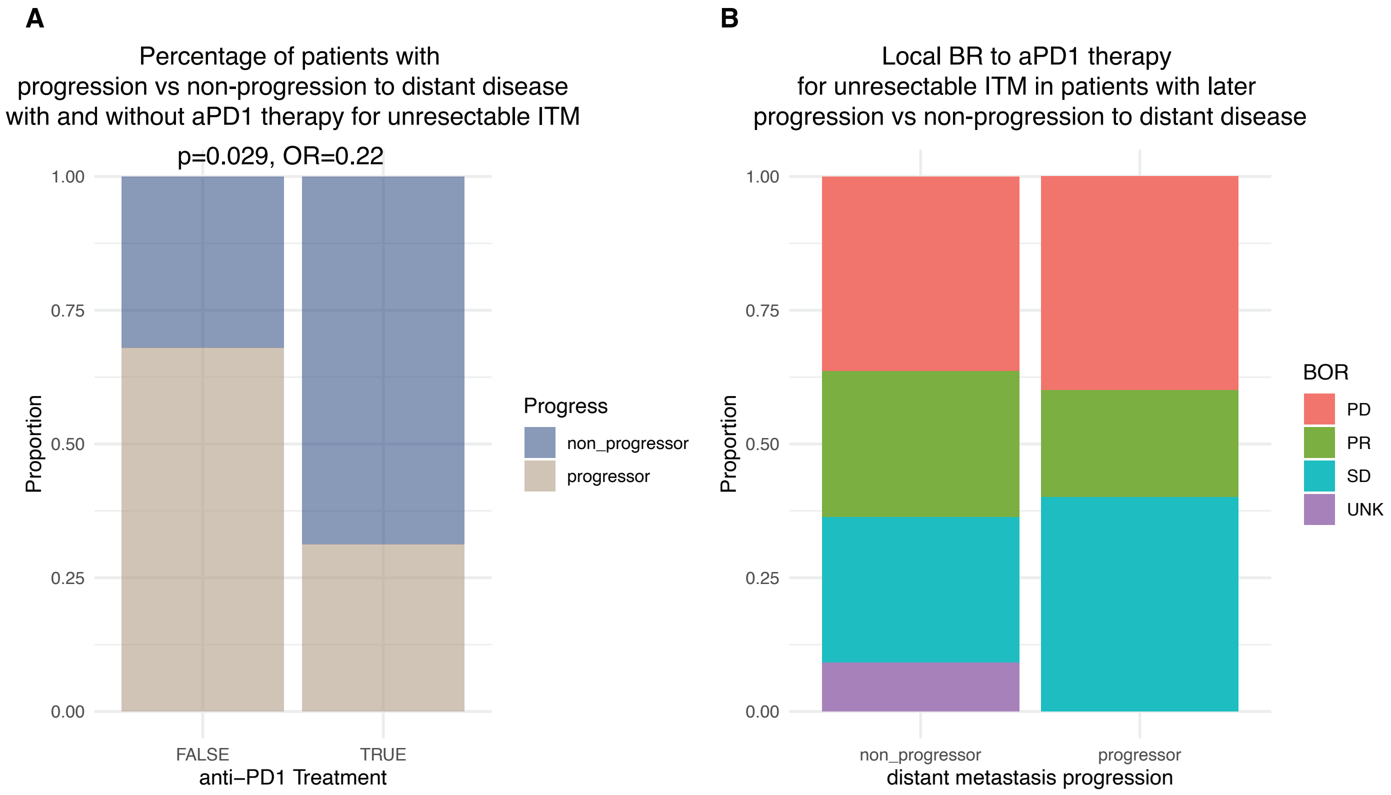
A
Percentage of patients with
progression vs non-progression to distant disease
with and without aPD1 therapy for unresectable ITM
p=0.029, OR=0.22
1.00
0.75
0.50
0.25
0.00
Proportion
FALSE
TRUE
anti−PD1 Treatment
Progress
non_progressor
progressor
B
Local BR to aPD1 therapy
for unresectable ITM in patients with later
progression vs non-progression to distant disease
1.00
0.75
0.50
0.25
0.00
Proportion
non_progressor
progressor
distant metastasis progression
BOR
PD
PR
SD
UNK

**Supplementary Figure 8. Investigating the role of the treatment with antiPD1 in ITM.**
**A.** Percentage of patients that have been treated or not with antiPD1 and the respective proportion of Progressors and non progressors in the two groups. **B.** Subset of patients treated with antiPD1 (n=16), stacked barplot showing the proportion of patients with partial response, stable disease and progressive disease.

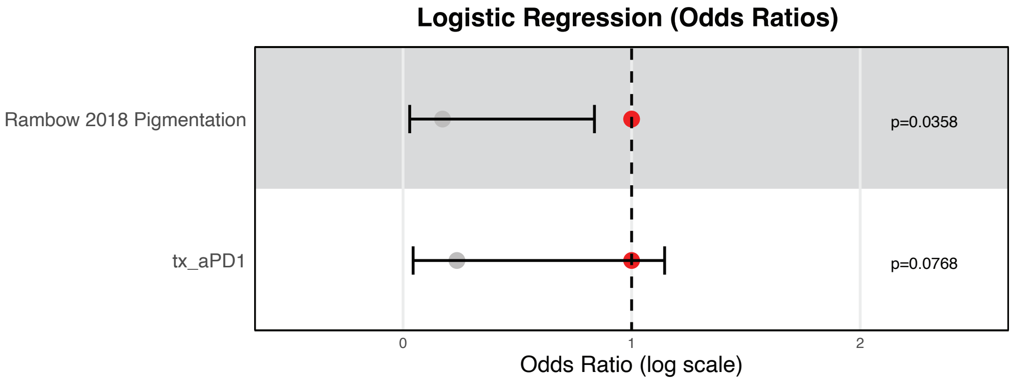

Logistic Regression (Odds Ratios)
Rambow 2018 Pigmentation
tx_aPD1
p=0.0358
p=0.0768
0
1
2
Odds Ratio (log scale)

**Supplementary Figure 9. Evaluation of the Pigmentation signature together with the antiPD1 treatment.**

**A.** Logistic regression model demonstrating that the *Rambow 2018 Pigmentation* signature and the treatment with antiPD1 are independent protective factors of disease progression. The pigmentation signature shows a protective effect with a significant odds ratio ($p$ = 0.0358), while the treatment with antiPD1 is also associated with a lower risk of progression but does not reach statistical significance ($p$ = 0.0768).

**A**

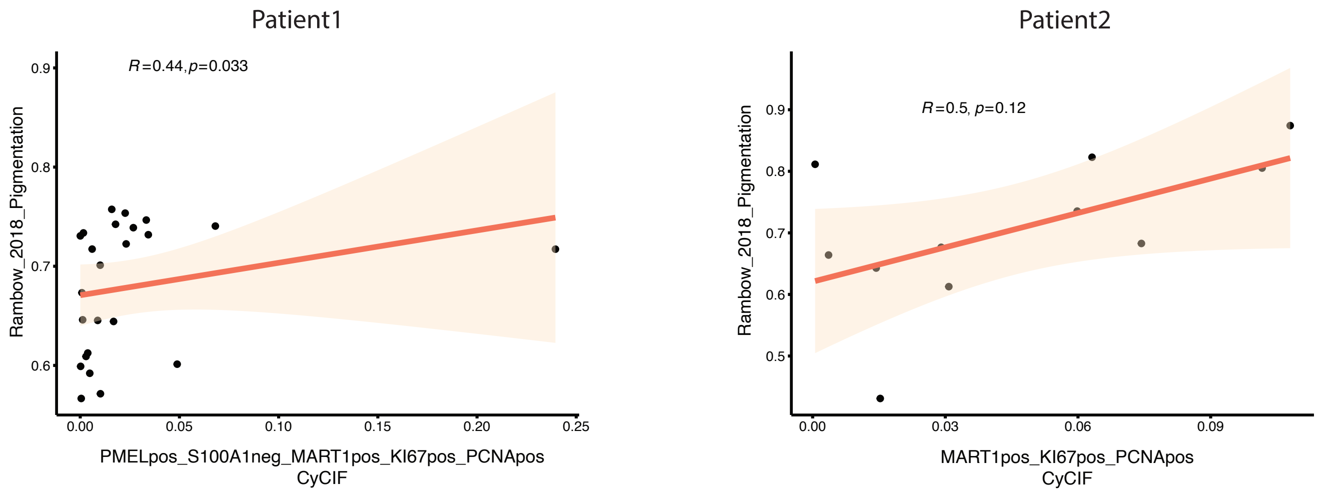


**B**

H&E from high Rambow_2018_Pigmentation signatures patients

H&E from low Rambow_2018_Pigmentation signatures patients

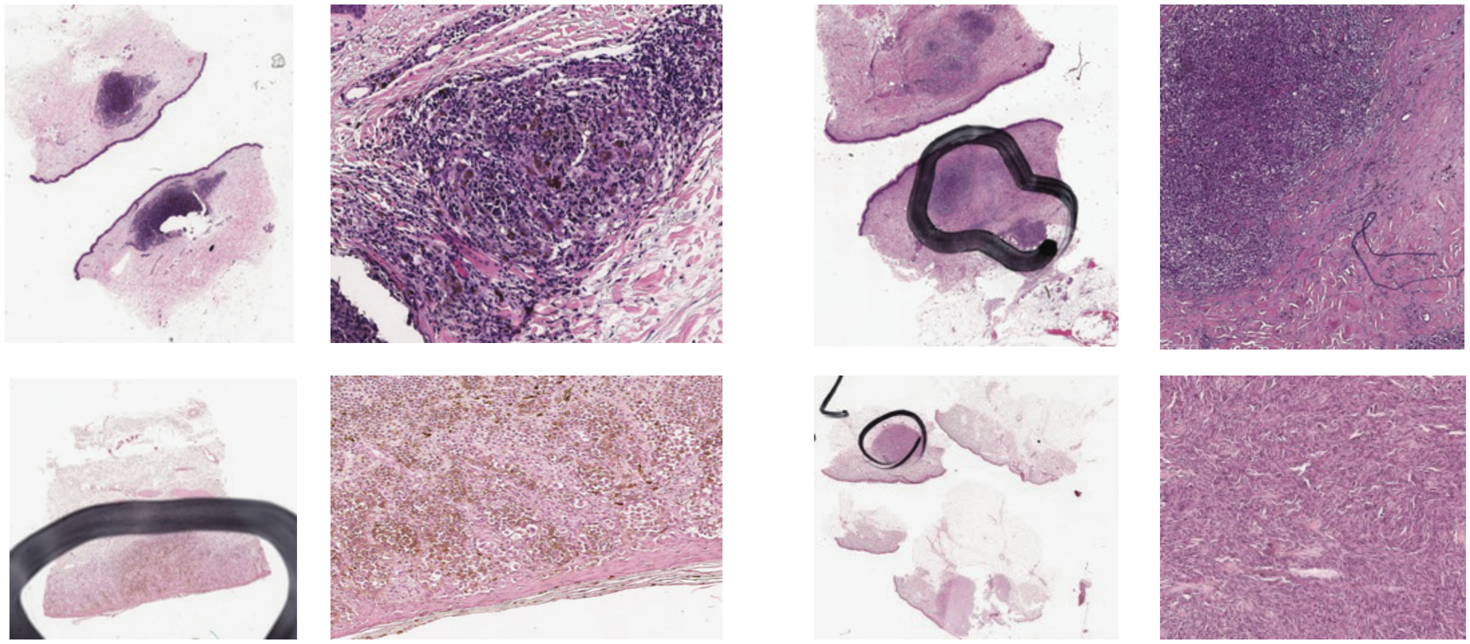

**Supplementary Figure 10. Signature scoring from CyCIF and H&E.**
**A.** Evaluating CyCIF melanocytic markers and their correlation with the pigmentation signature in patient1 (left) and patient2 (right). **B.** Comparison of the H&E of two samples with high pigmentation signature and two patients with low pigmentation signature. clear pigments visible from H&E evaluation from the patients with high pigmentation signature (estimated from bulk RNAseq)
